# The Temporal Stability of Visual Cortical Processing in Humans Depends on Early Experience

**DOI:** 10.1101/2025.03.25.645249

**Authors:** Suddha Sourav, Max Emanuel Feucht, Ramesh Kekunnaya, Brigitte Röder

## Abstract

Proper timing is essential for effective neural processing. Yet, how early postnatal vision shapes the temporal stability of human visual cortical processing remains unknown. Here, using electroencephalography, we examined cortical timing properties in individuals who were born pattern vision blind due to congenital cataracts, but surgically recovered sight. While sight-recovery individuals exhibited an attenuated cortical oscillatory phase coherence (i.e., higher temporal variability) during visual processing, their oscillatory strength was unimpaired. Moreover, phase coherence information, but not activation strength, allowed the classification of sight-recovery from control individuals. Finally, exchanging phase information between sight-recovery and control individuals indicated oscillatory timing impairments as the source of group differences in higher-order visual cortical processing. Neural timing impairments were specific for reversed congenital blindness, that is, were not observed in individuals with reversed developmental (late-onset) cataracts. These results suggest that the development of intricately temporally orchestrated visual cortical processing in humans requires early visual experience.

**Significance Statement:** Neural circuit functioning requires an efficient temporal orchestration. The present work investigated how the temporal stability of visual cortical dynamics depends on adequate experience. In rare individuals who were born with pattern vision blindness but later recovered sight, a marked increase in the temporal variability of oscillatory brain activity during visual processing was observed while oscillatory strength was surprisingly unimpaired. Therefore, the emergence of precisely timed visual processing in the human brain seems to crucially depend on early visual experience. We speculate that impaired timing of neural processing, cascading throughout the visual cortical hierarchy, might be a major source of the incomplete visual recovery in individuals with treated congenital blindness.

## Introduction

The typically developed brain exhibits a remarkable stability of sensory processing on a trial-by-trial basis, which may appear surprising in light of considerable spontaneous activity and diverse sources of biological noise in the brain (Rabinovich et al., 2008; Oby et al., 2025; Trägenap et al., 2025). Although numerous studies have investigated changes to visual cortical activation strength after sight loss (Wanet-Defalque et al., 1988; Rösler et al., 1993; Amedi et al., 2003), and subsequently, after sight recovery (Mitchell and Maurer, 2022; Röder and Kekunnaya, 2022), the neural mechanisms preventing complete behavioral recovery are yet unknown. In fact, humans who had suffered from a period of congenital pattern visual blindness due to the presence of total, bilateral congenital cataracts (CC), suffer from persistent visual impairments following sight restoration, even after extended recovery periods. These deficits include a permanently reduced visual acuity and stereoscopic ability (Tytla et al., 1993; Birch et al., 2009), atypical mid-to high-level visual feature processing (Putzar et al., 2007; McKyton et al., 2015), and altered face processing (Röder et al., 2013; De Heering and Maurer, 2014). In contrast, a substantially higher recovery of visual functions has been observed if the onset of visual loss was later (instead of congenital), e.g., through developmental cataracts (Röder et al., 2013; Bottari et al., 2018; Sourav et al., 2020). These results indicate the presence of *sensitive periods*, during which the effect of experience on the development of visual cortical circuits and the associated functions is especially strong (Knudsen, 2004; Maurer and Lewis, 2013; Röder et al., 2021). However, to date, the temporal stability of visual cortical processing as a sensitive period mechanism has not been studied in humans.

On the one hand, functional magnetic resonance imaging (fMRI) methods, indispensable for deriving spatially well-resolved activation maps in sight-recovery individuals, cannot resolve the fast, millisecond-level cortical temporal dynamics. On the other hand, electro/magnetoencephalographic (EEG/MEG) research methods have emphasized event-related (i.e., trial-averaged) indices of cortical activation (Makeig et al., 2004), discarding the trial-by-trial temporal variation information necessary to investigate the stability of visual cortical processing (Bottari et al., 2016; Ossandón et al., 2023; Sourav et al., 2024). For example, an attenuated event-related potential (ERP) in CC individuals, that is, a scalp-recorded marker of the summed electric field potentials in the brain, might result from a generally reduced cortical activation across trials, or equivalently, from a lack of temporal coherence across individual trials with unimpaired overall activation strength. At the level of the neural generators, these two hypothetical extremes represent fundamentally different (and to date not disentangled) effects of a period of transient visual deprivation, i.e., a generally attenuated cortical activity vs. compromised temporal stability of visual cortical processing.

In two experiments, we here investigated the temporal stability of visual cortical processing indexed by oscillatory phase coherence in sight recovery individuals with a history of transient congenital blindness. In addition, we compared cortical activation strength in CC individuals to normally sighted control individuals. First, time-frequency decomposition using wavelet transforms was employed to calculate the activation magnitude, operationalized by the post-stimulus power change of cortical oscillations compared to a pre-stimulus baseline, as well as the inter-trial oscillatory phase coherence to assess the temporal stability of cortical visual processing. These two markers were subsequently compared between individuals with reversed congenital cataracts and normally sighted controls, and to ascertain the presence of an early sensitive period, additionally between individuals with reversed developmental (childhood) cataracts and normally sighted controls. Next, we used a classification approach that attempted to identify CC individuals based on power changes relative to pre-stimulus baseline, as well as based on inter-trial phase coherence information in the time-frequency representations. Further, we derived what we term *“mutual magnitude-phase transplants”,* a novel method that we here introduce, which allowed us to exchange the per-trial activation strength and timing information between a cataract reversal individual and their matched normally sighted control participant. Previously, we have reported a robust and replicable attenuation of the earliest electrophysiological marker of higher-order (extrastriate) visual cortical processing, the P1 wave (∼ 150 ms; Di Russo et al., 2002; Miller et al., 2015), in CC individuals (Sourav et al., 2020). The method of mutual magnitude-phase transplants allowed us to investigate the relative contribution of activation strength vs. timing information to the extrastriate cortical changes observed after a period of transient congenital visual deprivation. Finally, minimum-norm source estimation was performed to verify the early extrastriate cortical sources of the P1 wave.

## Materials and Methods

### Participants

Datasets originating from two different experiments, termed experiment 1 and experiment 2, were utilized. Both experiments included a group of congenital cataract reversal individuals (CC) as well as a group of developmental cataract reversal individuals (DC), who had undergone the same surgical procedures. In addition, normally sighted control participants with a typical developmental history were recruited, and each normally sighted control participant was matched to a cataract reversal individual for age, sex, and handedness.

Thirteen CC individuals took part in experiment 1, of whom the data of 1 participant were not analyzed here due to a very short visual deprivation period (1 month). The remaining CC participants, whose data entered further analysis (*n* = 12, see Table 1 for participant details), had a mean age of 18.67 years (range = 10 – 37 years), 2 were female (and 10 were male), and 1 was left-handed. In addition, 16 DC individuals took part in experiment 1, of whom the data of 3 individuals were not analyzed respectively due to a diagnosis of seizure and developmental delay, the participant’s unwillingness to continue the experiment shortly after the start and the inability of the participant to look straight ahead. The data of 13 DC individuals (Table 2) entered further analysis (mean age = 16 years, range = 11 – 24 years, 5 females and 8 males, all right-handed). The CC individuals had received surgical treatment after a mean of 41.33 months of blindness (range 4 – 213 months). The average age at treatment for DC individuals was 117.38 months (range 24 – 208 months). The visual acuity (VA) of the CC individuals (geometric mean, decimal = 0.23, range = 0.05 – 0.5) was significantly lower compared to that of the DC individuals (geometric mean, decimal = 0.74, range = 0.44 – 1.00), as revealed by a one-tailed *t*-test after converting the VAs to LogMAR units, *t*(14.531) = 5.642, *p* < .001. The mean age of the typically sighted control individuals matched to the CC individuals (MCC group) was 18.08 years (range = 10 – 34 years, 2 female and 10 male, 1 left-handed), and 16.85 years for the typically sighted control individuals matched to the DC individuals (MDC group, range = 10 – 25 years, 5 female and 8 male, all right-handed).

**Table 1.**
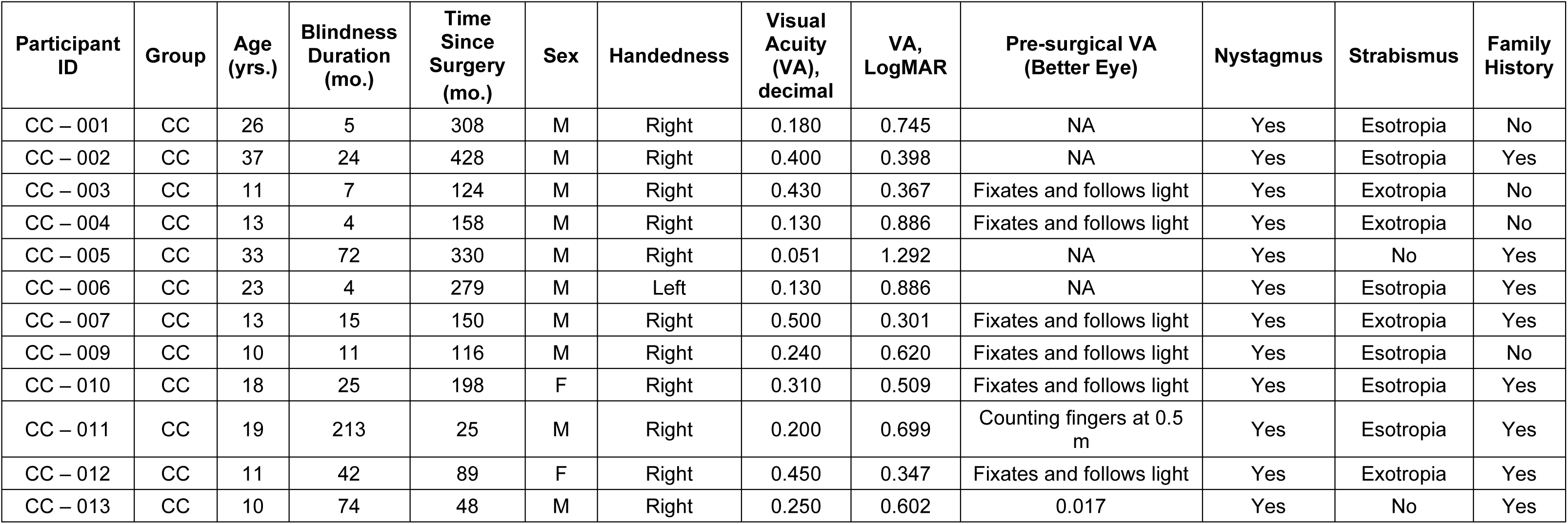
Participant characteristics for the sight recovery individuals with a history of dense bilateral congenital cataracts and subsequent surgery in experiment 1. The table is based on Sourav et al. (2020).

| Participant ID | Group | Age (yrs.) | Blindness Duration (mo.) | Time Since Surgery (mo.) | Sex | Handedness | Visual Acuity (VA), decimal | VA, LogMAR | Pre-surgical VA (Better Eye) | Nystagmus | Strabismus | Family History |
| --- | --- | --- | --- | --- | --- | --- | --- | --- | --- | --- | --- | --- |
| CC – 001 | CC | 26 | 5 | 308 | M | Right | 0.180 | 0.745 | NA | Yes | Esotropia | No |
| CC – 002 | CC | 37 | 24 | 428 | M | Right | 0.400 | 0.398 | NA | Yes | Esotropia | Yes |
| CC – 003 | CC | 11 | 7 | 124 | M | Right | 0.430 | 0.367 | Fixates and follows light | Yes | Exotropia | No |
| CC – 004 | CC | 13 | 4 | 158 | M | Right | 0.130 | 0.886 | Fixates and follows light | Yes | Exotropia | No |
| CC – 005 | CC | 33 | 72 | 330 | M | Right | 0.051 | 1.292 | NA | Yes | No | Yes |
| CC – 006 | CC | 23 | 4 | 279 | M | Left | 0.130 | 0.886 | NA | Yes | Esotropia | Yes |
| CC – 007 | CC | 13 | 15 | 150 | M | Right | 0.500 | 0.301 | Fixates and follows light | Yes | Exotropia | Yes |
| CC – 009 | CC | 10 | 11 | 116 | M | Right | 0.240 | 0.620 | Fixates and follows light | Yes | Esotropia | No |
| CC – 010 | CC | 18 | 25 | 198 | F | Right | 0.310 | 0.509 | Fixates and follows light | Yes | Esotropia | Yes |
| CC – 011 | CC | 19 | 213 | 25 | M | Right | 0.200 | 0.699 | Counting fingers at 0.5 m | Yes | Esotropia | Yes |
| CC – 012 | CC | 11 | 42 | 89 | F | Right | 0.450 | 0.347 | Fixates and follows light | Yes | Exotropia | Yes |
| CC – 013 | CC | 10 | 74 | 48 | M | Right | 0.250 | 0.602 | 0.017 | Yes | No | Yes |

**Table 2.** Participant characteristics for the sight recovery individuals with a history of bilateral developmental cataracts and subsequent surgery (DC) in experiment 1. The table is based on Sourav et al. (2020).

| Participant ID | Group | Age (yrs.) | Age at Surgery (yrs.) | Time Since Surgery (mo.) | Sex | Handedness | Visual Acuity (VA), decimal | VA, LogMAR | Pre-surgical VA (Better Eye) | Nystagmus | Strabismus | Family History |
| --- | --- | --- | --- | --- | --- | --- | --- | --- | --- | --- | --- | --- |
| DC – 001 | DC | 16 | 12 | 53 | M | Right | 1.000 | 0.000 | Counting fingers at 0.5 m | No | No | No |
| DC – 002 | DC | 19 | 14 | 55 | M | Right | 1.000 | 0.000 | 0.200 | No | No | No |
| DC – 003 | DC | 12 | 6 | 63 | F | Right | 1.000 | 0.000 | 0.500 | No | No | No |
| DC – 004 | DC | 11 | 8 | 39 | F | Right | 0.530 | 0.276 | 0.159 | No | No | No |
| DC – 005 | DC | 13 | 8 | 68 | M | Right | 0.710 | 0.149 | 0.100 | No | Exotropia | No |
| DC – 006 | DC | 12 | 8 | 51 | M | Right | 0.600 | 0.222 | 0.400 | No | No | No |
| DC – 007 | DC | 12 | 6 | 77 | M | Right | 1.000 | 0.000 | 0.250 | No | No | Yes |
| DC – 008 | DC | 18 | 12 | 74 | F | Right | 0.790 | 0.102 | Sees hand movement | No | No | No |
| DC – 009 | DC | 16 | 10 | 67 | M | Right | 0.740 | 0.131 | 0.400 | No | No | No |
| DC – 010 | DC | 24 | 2 | 264 | F | Right | 0.440 | 0.357 | Fixates and follows light | No | Exotropia | No |
| DC – 011 | DC | 16 | 7 | 110 | F | Right | 0.530 | 0.276 | Counting fingers @ 1 m | No | Exotropia | Yes |
| DC – 012 | DC | 20 | 12 | 97 | M | Right | 0.880 | 0.056 | 0.400 | No | No | No |
| DC – 013 | DC | 19 | 17 | 17 | M | Right | 0.730 | 0.137 | 0.200 | No | No | No |

In experiment 2, 15 participants with a history of bilateral, dense congenital cataracts took part. The data of 2 of these participants were rejected because in one case, the participant was born with incomplete cataract and thus did not suffer from complete pattern visual blindness since birth, and in another case, because of a very short visual deprivation period (1 month). Thus, the data of 13 CC individuals entered further analysis (Table 3, mean age = 17.69 years, range 6 – 39 years, 3 female and 10 male, 1 left-handed). In addition, 16 DC individuals took part in experiment 2. We did not analyze the data of 1 participant due to a history of seizures and developmental delay. The data of 15 DC individuals entered further analysis (Table 4, mean age = 14.47 years, range 9 – 24 years, 4 female and 11 male, 1 left-handed).

**Table 3.** Participant characteristics for the sight recovery individuals with a history of dense bilateral congenital cataracts and subsequent surgery (CC) in experiment 2 (Please note that age of the same participant might vary from. **Table 1 due to the experiments being run at different time points). The table is based on Sourav et al. (2020)**.

| Participant ID | Group | Age (yrs.) | Blindness Duration (mo.) | Time Since Surgery (mo.) | Sex | Handedness | Visual Acuity (VA), decimal | VA, LogMAR | Pre-surgical VA (Better Eye) | Nystagmus | Strabismus | Family History |
| --- | --- | --- | --- | --- | --- | --- | --- | --- | --- | --- | --- | --- |
| CC – 001 | CC | 26 | 5 | 308 | M | Right | 0.180 | 0.745 | NA | Yes | Esotropia | No |
| CC – 005 | CC | 33 | 72 | 330 | M | Right | 0.051 | 1.292 | NA | Yes | No | Yes |
| CC – 014 | CC | 6 | 5 | 70 | M | Right | 0.240 | 0.620 | Not fixating/following light | Yes | No | No |
| CC – 002 | CC | 39 | 24 | 444 | M | Right | 0.400 | 0.398 | NA | Yes | Esotropia | Yes |
| CC – 004 | CC | 14 | 4 | 175 | M | Right | 0.130 | 0.886 | Fixates and follows light | Yes | Exotropia | No |
| CC – 006 | CC | 24 | 4 | 294 | M | Left | 0.130 | 0.886 | NA | Yes | Esotropia | Yes |
| CC – 007 | CC | 15 | 15 | 166 | M | Right | 0.500 | 0.301 | Fixates and follows light | Yes | Exotropia | Yes |
| CC – 009 | CC | 11 | 11 | 129 | M | Right | 0.240 | 0.620 | Fixates and follows light | Yes | Esotropia | No |
| CC – 015 | CC | 8 | 48 | 48 | F | Right | 0.150 | 0.824 | Perception of light | Yes | No | Yes |
| CC – 013 | CC | 11 | 74 | 61 | M | Right | 0.250 | 0.602 | 0.017 | Yes | No | Yes |
| CC – 012 | CC | 11 | 42 | 90 | F | Right | 0.450 | 0.347 | Fixates and follows light | Yes | Exotropia | Yes |
| CC – 011 | CC | 21 | 213 | 40 | M | Right | 0.200 | 0.699 | Counting fingers at 0.5 m | Yes | Esotropia | Yes |
| CC – 016 | CC | 11 | 72 | 68 | F | Right | 0.260 | 0.585 | Fixates and follows light | Yes | No | No |

**Table 4.** Participant characteristics for the sight recovery individuals with a history of bilateral developmental cataracts and subsequent surgery (DC) in experiment 2 (Please note that age of the same participant might vary from. **Table 2 due to the experiments being run at different time points). The table is based on Sourav et al. (2020)**.

| Participant ID | Group | Age (yrs.) | Age at Surgery (yrs.) | Time Since Surgery (mo.) | Sex | Handedness | Visual Acuity (VA), decimal | VA, LogMAR | Pre-surgical VA (Better Eye) | Nystagmus | Strabismus | Family History |
| --- | --- | --- | --- | --- | --- | --- | --- | --- | --- | --- | --- | --- |
| DC – 002 | DC | 19 | 14 | 55 | M | Right | 1.000 | 0.000 | 0.200 | No | No | No |
| DC – 011 | DC | 14 | 7 | 80 | F | Right | 0.530 | 0.276 | Finger counting at 1 m | No | Exotropia | Yes |
| DC – 003 | DC | 13 | 6 | 74 | F | Right | 1.000 | 0.000 | 0.500 | No | No | No |
| DC – 005 | DC | 14 | 8 | 80 | M | Right | 0.710 | 0.149 | 0.100 | No | Exotropia | No |
| DC – 007 | DC | 12 | 6 | 77 | M | Right | 1.000 | 0.000 | 0.250 | No | No | Yes |
| DC – 014 | DC | 12 | 7 | 66 | M | Right | 0.230 | 0.638 | 0.125 | No | No | No |
| DC – 015 | DC | 12 | 7 | 67 | M | Right | 0.460 | 0.337 | 0.100 | No | No | No |
| DC – 009 | DC | 16 | 10 | 67 | M | Right | 0.740 | 0.131 | 0.400 | No | No | No |
| DC – 010 | DC | 24 | 2 | 264 | F | Right | 0.440 | 0.357 | Fixates and follows light | No | Exotropia | No |
| DC – 008 | DC | 18 | 12 | 74 | F | Right | 0.790 | 0.102 | Perceives hand movement | No | No | No |
| DC – 016 | DC | 17 | 12 | 60 | M | Right | 0.310 | 0.509 | 0.160 | No | No | Yes |
| DC – 017 | DC | 9 | 2 | 86 | M | Right | 0.930 | 0.032 | 0.154 | No | No | No |
| DC – 018 | DC | 16 | 6 | 113 | M | Right | 0.520 | 0.284 | Finger counting at 1 m | No | No | No |
| DC – 019 | DC | 10 | 3 | 76 | M | Left | 0.730 | 0.137 | 0.065 | No | No | Yes |
| DC – 020 | DC | 11 | 1 | 115 | M | Right | 0.667 | 0.176 | 0.154 | No | No | No |

The CC individuals in experiment 2 whose data were analyzed (*n* = 13) had been treated for cataract after a mean of 45.31 months of blindness (range 4 – 213 mo.), and the DC individuals whose data were analyzed had been treated for cataract with a mean age of 88.33 months (range 23 – 173 mo.). Once again, the CC individuals exhibited a generally lower mean geometric visual acuity (0.21, range = 0.051 – 0.5) than the DC individuals (0.618, range = 0.23 – 1.00; one-tailed t-test, *t*(21.532) = 5.297, p < .001). The MCC group in experiment 2 had a mean age of 17 years (range = 7 – 38 years, 3 female and 10 male, 1 lefthanded), and the MDC group had a mean age of 14.93 years (range = 11 – 26 years, 4 female and 11 male, 1 lefthanded).

Ten CC individuals, 8 DC individuals and 5 normally sighted control individuals took part in both experiment 1 and experiment 2; 35.7% of cataract reversal individuals were therefore unique to experiment 2 (i.e., 3 CC and 7 DC individuals). All cataract reversal individuals from both experiments were recruited and tested at the L V Prasad Eye Institute, Hyderabad, India. Control individuals were tested at the University of Hamburg, Hamburg, Germany. Adult participants received a small monetary compensation for taking part in the study, whereas minors received a small gift. Costs for travel and accommodation were reimbursed. The study obtained approval from the local ethics commission of the faculty of Psychology and Human Movement at the University of Hamburg and the institutional ethics review board of LVPEI, and adhered to the ethical principles outlined in the Declaration of Helsinki (World Medical Association, 2013) .

### Experimental Design

In both experiments, participants were presented monochromatic square wave grating stimuli in front of a grey background, one at a time for 150 ms in one of the four visual field quadrants with a mean interstimulus interval of 1.85 s (1.5 s – 2.2 s). The stimuli were presented with a fixed eccentricity of 4 degrees from the center and subtended a visual angle of 2.5° as depicted in Figure 2 in Sourav et al. (2020). The stimuli were presented at a polar angle of 25° in the upper visual field and at a polar angle of −45° in the lower visual field. The gratings were either horizontally (standard stimuli, *P* = .80) or vertically oriented (target stimuli, *P* = .20), with a spatial frequency of 2 cycles/degree. Participants were instructed to attend to a central fixation cross and to indicate when a rare target stimulus appeared in any of the locations by pressing a foot pedal in experiment 1 and by clicking a computer mouse in experiment 2. Sight restored cataract individuals were tested using a Dell IN2030 monitor at LVPEI, Hyderabad, and typically sighted control individuals were tested on a Samsung P2370 monitor at the University of Hamburg. Both monitors had a refresh rate of 60 Hz and a nominal luminance of 250 cdm^-2^. Participants sat at a distance of 45 cm to the monitor in experiment 1, and at a distance of 65 cm in experiment 2. The angular measurements described above were identical in both experiments. Participants were presented additional stimuli in both experiments which are not reported here.

**Figure 1.**
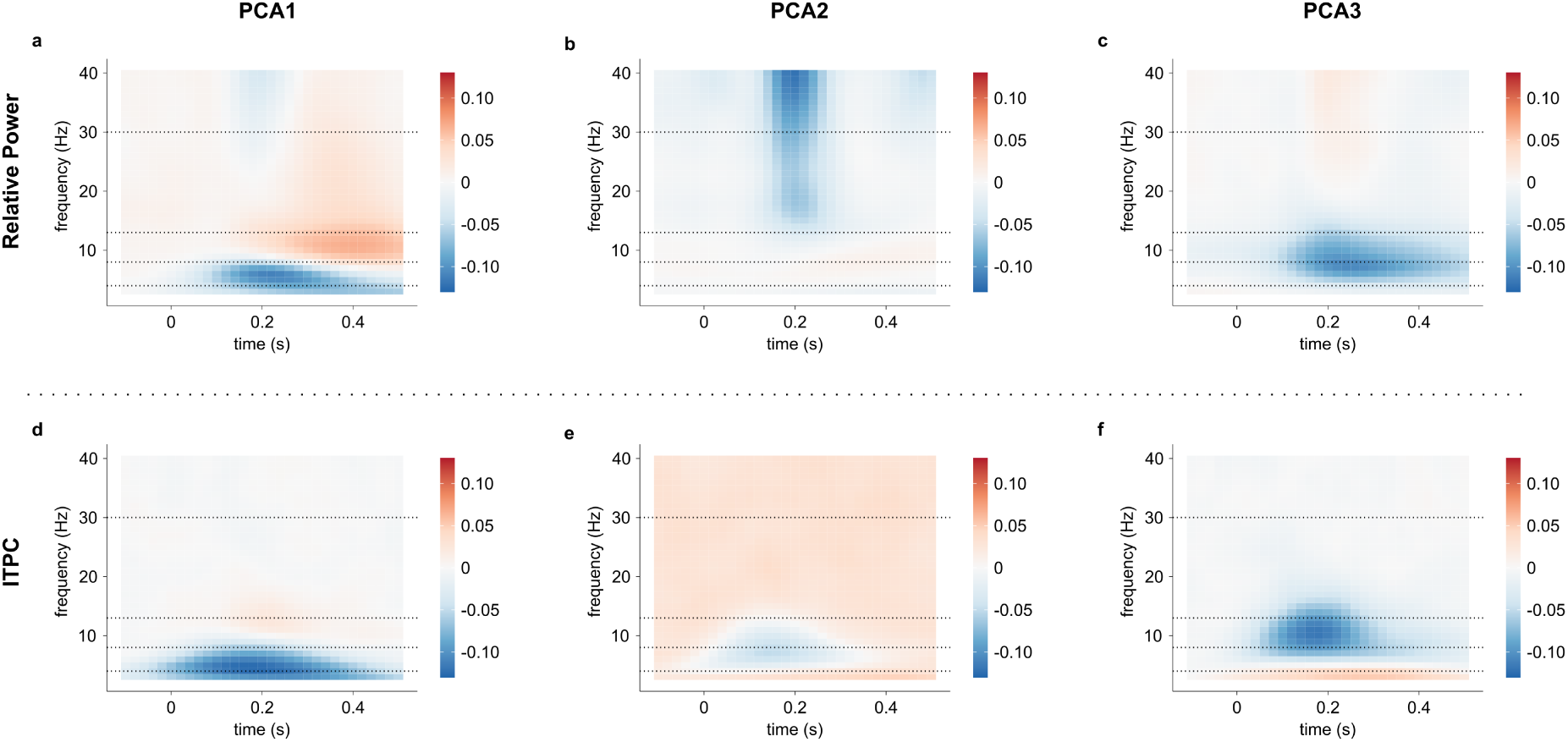
Principal component loadings of relative power change and of ITPC. **a – c**. The first three principal components of the relative power change compared to baseline (*ΔP*rel), summed across all posterior electrodes for each time-frequency feature (unitless). **d – e**. The first three principal components of inter-trial phase coherence (ITPC), summed across all posterior electrodes for each time-frequency feature (unitless). Each pixel in each plot represents one time-frequency feature. Summation over all electrodes is performed to highlight overall positive and negative loading clusters for each component.

**Figure 2.**
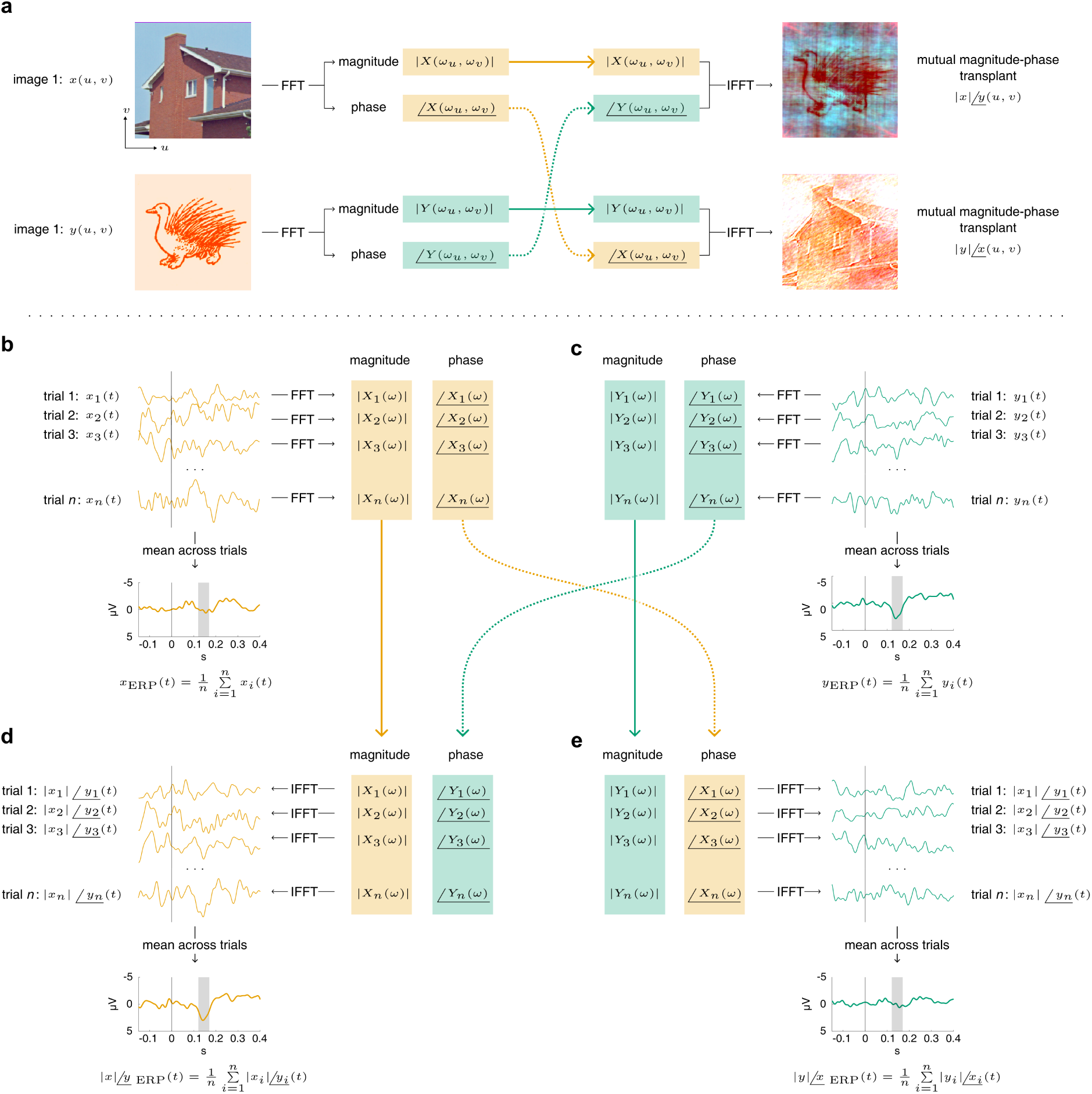
Derivation of mutual magnitude-phase transplants (MuTx). **a** An image processing example motivating the MuTx approach. Two-dimensional fast Fourier transform (FFT) of signals, here public-domain images of a house (Weber, 2018) and an imaginary animal (Roy, 1946), decomposes each image into a set of sinusoidal spatial oscillations (here, in two spatial direction *u* and *v*). Each oscillation (i.e. each member of the set of sinusoidal oscillations derived by FFT) have a magnitude, indicating the strength of oscillation, as well as a phase, indicating its relative position. As an experiment, the magnitude and phases of the two images can be exchanged. After this interchange, inverse fast Fourier transforms (IFFT) yield the mutual magnitude-phase transplants (MuTx). It can be appreciated that for images, most of the shape information seems to be contained in the signal phase, whereas color information seems to rely more on the signal magnitude. **b – c** Derivation of magnitude and phase information via one-dimensional FFT of the individual trials of the EEG epochs respectively in a CC individual and their matched normally sighted control participant (MCC), with the mean across trials (unaltered event related potentials, ERPs) displayed. **d – e** After exchanging the magnitude and phase spectra, the inverse FFT yields two MuTx waves with the magnitude of the CC individual but the phase of the MCC individuals (|𝐶𝐶|∠𝑀𝐶𝐶) and vice versa (|𝑀𝐶𝐶|∠𝐶𝐶). It can be appreciated that phase information seems to drive most of the P1 magintude difference (grey bar, 120 – 170 ms post-stimulus) between the CC and the MCC participant.

For stimulus presentation we used the the PsychoPy framework (v1.83, Peirce, 2009).

### EEG: Data Acquisition and Preprocessing

EEG data were acquired continuously during the whole experiment using 32 Ag/AgCl electrodes with a left earlobe online reference. Brain activity was recorded at the 10-10 locations FP1, FP2, F7, F3, Fz, F4, F8, FC5, FC1, FCz, FC2, FC6, T7, C3, Cz, C4, T8, TP9, CP5, CP1, CP2, CP6, TP10, P7, P3, Pz, P4, P8, O1, O2, F9, and F10, with an online high-pass filter with a cutoff frequency of 0.016 Hz, a low-pass filter with a cutoff frequency of 250 Hz, and a sampling rate of 1 kHz. To eliminate electrical line noise, the data were notch-filtered offline at 50 Hz and, if apparent in the frequency spectrum, at the corresponding harmonics. Subsequently, the voltages were average referenced and biological artifacts such as blinks, eye movements, muscle activity or heartbeats were removed from the signal with an independent component analysis (ICA), using EEGLAB for MATLAB (EEGLAB, version 11.5.4b, Delorme and Makeig, 2004; MATLAB version 2012b; MathWorks, Natick, MA). To prevent confounds due to eye movements or blinks during stimulus presentation, we excluded stimulus epochs that contained blinks or eye movements within the time window of −25 ms to 175 ms relative to the stimulus, as previously reported (Sourav et al., 2020, 2024). The data were then low pass filtered at 40 Hz and EEG epochs were generated from -1.7s before to 1.7 s after the stimulus onset. For increasing the statistical power, data from trials with a left hemifield stimulation were remapped; at each electrode, the data were swapped with the data of the electrode in its mirrored location with respect to the nasion-inion axis, ignoring the midline electrodes (see Sourav et al., 2020). Additionally, to prevent artifacts related to motor activation, epochs containing a motor response within ±500 ms of the stimulus onset were rejected.

Only EEG trials/ERPs generated by the frequent, standard stimuli were used for further analysis. Moreover, as extrastriate-generated P1 wave amplitudes elicited by upper visual field stimulation are larger than those elicited by lower visual field stimulations in the present paradigm, we only considered visual ERPs for upper visual field stimulation for the subsequent analyses, as in Sourav et al. (2020).

### Data Reuse

The dataset in the current study has been derived from Sourav et al. (2020). The only difference is the exclusion of one participant with a very short duration of visual deprivation prior to the commencement of data analysis (1 month in participant CC – 008; see supplemental materials for Sourav et al., 2020). Parts of the dataset have been included in two other publications (Sourav et al., 2018, 2024).

### Time-Frequency Analysis

Time-frequency representations of the EEG data were derived by wavelet transformation of individual trials at each electrode using the *FieldTrip* software in MATLAB (Oostenveld et al., 2011). Signals at each electrode were convolved with complex Morlet wavelets which have a Gaussian shape over time (with standard deviation σ_t_) as well as over frequency (with standard deviation σ_f_), centered around a time t_0_ and a frequency f_0_ (Tallon-Baudry et al., 1999). Non-constant wavelets were applied with a ratio f_0_/σ_f_ = 4 for the lowest frequency and increasing linearly up to 8.5 for the highest, identical to Bottari et al. (2016, 2018). Thereafter, the relative power change compared to baseline (*ΔP*_rel_), and the inter-trial phase coherence (ITPC) across trials were computed at each frequency from 2 to 40 Hz with a step size of 1 Hz, and from 700 ms before to 700 ms after stimulus onset in steps of 20 ms. *ΔP*_rel_ was derived by normalizing the wavelet power with respect to a baseline interval from -700 ms to -300 ms, i.e., *ΔP*(t,f)_rel_ = [P(t,f) – P_baseline_(f)]/P_baseline_(f), see Bottari et al. (2016, 2018). The ITPC is a measure of oscillatory phase synchronization across trials at a given timepoint and frequency (Delorme & Makeig, 2004) and was derived by calculating the magnitude of the complex average of the normalized phase directions. The ITPC can take on values in the interval [0, 1], where ITPC = 1 indicates a perfect synchronization across trials and ITPC = 0 indicates complete desynchronization. Importantly, the ITPC is a biased but consistent statistical measure, e.g., for uniformly random, desynchronized phase values with a finite number of trials, the ITPC is expected to be positive (i.e., is overestimated), but approaches 0 as the number of trials is increased (Aydore et al., 2013). To remove this bias of the ITPC estimates for group comparisons, and to derive mutual magnitude-phase transplants as described in a following section, we used an equal number of EEG epochs for each cataract reversal participant and their control participant.

### Cluster Based Permutation Tests

The *ΔP*_rel_ and ITPC values were compared between each cataract reversal group and their sighted controls (i.e., CC vs. MCC; DC vs. MDC) using two-tailed cluster-based permutation tests. The tests were performed with the *FieldTrip* framework (Oostenveld et al., 2011) running on MATLAB (R2022a, Natick, MA), over the post-stimulus time-frequency plane (*t*: 0 – 700 ms; *f*: 2- 40 Hz) and over the 13 electrodes posterior to the midline, where the P1 wave is most prominent. The cluster-based permutation tests were based on 10,000 randomizations and used weighted cluster mass as the test metric.

After visual stimulus presentation, a task-related decrease of *ΔP*_rel_ is typically observed in the alpha (8 – 12 Hz) frequency range (“alpha blocking”, Klimesch, 2012). Based on a previous report by Bottari et al. (2016) that task-related post-stimulus alpha blocking might differ in the CC group, we additionally compared *ΔP*_rel_ in each sight recovery group to their control group (i.e., CC vs. MCC; DC vs. MDC) with cluster-based permutation tests, with the same parameters as above, in the “alpha blocking” time-frequency window (time: 0.3 – 0.7 s, frequency: 8 – 12 Hz) .

### Classification Based on Time-Frequency Features

In addition to statistical tests for group differences in time-frequency representations, we tested group discriminability of CC vs. Non-CC (i.e., DC and MCC) and CC vs. DC individuals in a classification paradigm based on the time-frequency features, separately for *ΔP*_rel_ and ITPC. Classification was performed based on oscillatory activity from 2 to 40 Hz in steps of 1 Hz (i.e., in 39 frequency bins) and from 100 ms before stimulus onset to 500 ms after stimulus onset, with 20 ms step size (i.e., 31 temporal bins) at all 13 posterior electrodes, yielding 15717 classification features for both *ΔP*_rel_ and ITPC. To deal with the high dimensionality of the data, we employed principal component analysis (PCA) prior to classification (Buzzell et al., 2022). Using the principal component loadings, participants were then classified using linear support vector machines (SVM). Data from experiment 1 were used for deriving the principal components, for tuning the SVM hyperparameters, and for subsequent SVM training. In contrast, data from experiment 2 were utilized to validate the previously trained SVMs. The analysis was performed in R, version 4.0.3 (R Core Team, 2016).

### Dimension Reduction of the Time-Frequency Features

The *ΔP*_rel_ and ITPC values at each frequency and timepoint at the 13 posterior electrodes were used to derive principal components for *ΔP*_rel_ and ITPC separately. PCA was performed across all participants of experiment 1 (including the normally sighted control individuals), without centering or scaling the *ΔP*_rel_ and ITPC features, respectively. We chose the first three principal components for classification which explained 84.84% of the variance of ITPC and 49.21% of the variance of *ΔP*_rel_. Importantly, the PCA components do not necessarily reflect components with the highest *intergroup* variance, rather they reflect the highest *interindividual* variance regardless of the group membership. According to the suggestion by Buzzell et al. (2022), varimax rotation was first applied to the selected components before computing the single participants’ loadings at each electrode, yielding 39 *ΔP*_rel_ and 39 ITPC features per participant (13 electrodes × 3 principal component loadings). The varimax-rotated components are displayed in Fig. 1.

To assess the robustness of potential group differences, classification was additionally performed based on loadings of each of the first three components averaged over all posterior electrodes, yielding 3 classification features for both *ΔP*_rel_ and ITPC. The principal components were solely derived from data of experiment 1 and the same components were used for classification of the data from experiment 2.

### Support Vector Machine Configuration

Using the features derived from the PCA-based dimensionality reduction, the regularizing hyperparameter *c* of the SVM was tuned in a repeated cross-validation (CV) procedure, with a split of *k* = 5 (CC vs. non-CC classification) and *k* = 3 (CC vs. DC classification). The CV procedure was repeated 500 times, and the optimal *c* that was chosen most often of all repetitions was used to fit a final SVM to all data from experiment 1. For each feature class (i.e., *ΔP*_rel_ and ITPC), each classification type (CC vs. non-CC, and CC vs. DC), an SVM was trained once using all 39 component loadings from experiment 1 and the most often chosen optimal value of *c*, summing to 4 training procedures (2 feature classes, × 2 classification types). This procedure was repeated for the component loadings averaged over the posterior electrodes, yielding 4 additional training procedures each only using 3 classification features. For each SVM, a receiver operating characteristic (ROC) analysis was conducted to extract the optimal classification threshold by maximizing the *Youden’s J* parameter. To validate each trained classifier, the decision boundary derived from experiment 1 was used to classify the participants in experiment 2. ROC analyses were conducted for each classifier and the corresponding areas under the ROC curves (AUCs) were extracted. Subsequently, the significance of the AUCs was determined using the Mann-Whitney U-Statistic (Mason and Graham, 2002), and the resulting *p*-values were adjusted using the Benjamini and Hochberg procedure (Benjamini and Hochberg, 1995). The sensitivity and specificity of each classification procedure was obtained through the optimal thresholds derived from experiment 1.

### Mutual Magnitude-Phase Transplants

The discrete Fourier transform decomposes a signal (e.g., scalp-recorded voltage at an electrode) into a set (“spectrum”) of complex sinusoidal oscillations, each possessing a different frequency. A constituent oscillation has a magnitude, indicating the oscillation strength, as well as a phase, indicating its relative position in time or space (Grami, 2016). The magnitude and phase spectra of a signal generally carry distinct information about the signal. For example, in an image, most of the global color information is held in the magnitude spectrum of the color channels, whereas the phase spectrum contains most of the form information (Weinhaus, 2011). It is well known that swapping the magnitude and phase information between two images creates chimeric images that generally inherit the color palette from the magnitude spectrum and the form information from the phase spectrum (Fig. 2a). To probe the relative contribution of signal strength vs. coherent timing to the P1 wave reduction observed in CC individuals, we adapted this signal processing method for event-related potentials at the level of individual trials. Here we briefly describe this method, which we term *mutual magnitude-phase transplants* (MuTx) in the present context.

Event-related potentials (ERP) are derived by averaging across numerous trials (Fig. 2 b – c). For each CC individual and their matched normally sighted control participant (MCC), we derived the MuTx waves from a set balanced number of trials, as in all the other analyses reported in this article. First, for both individuals, we applied a Hann window to each individual trial and calculated the discrete Fourier transform with the FFT method in Matlab (R2022a, Natick, MA). Then, the magnitude and the phase spectra of the FFT was calculated, e.g.,

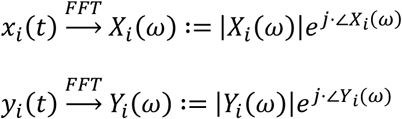

where |𝑋_𝑖_(𝜔)| and ∠𝑋_𝑖_(𝜔) are the magnitude and phase spectra of the 𝑖^𝑡ℎ^ trial of the CC individual, |𝑌_𝑖_(𝜔)| and ∠𝑌_𝑖_(𝜔) are the magnitude and phase spectra of the 𝑖^𝑡ℎ^ trial of the MCC individual, and 𝑗 is the imaginary unit, i.e. 𝑗^2^ = −1.

After this step, the phase of the CC and the MCC individuals were exchanged, and an inverse fast Fourier transform (IFFT) yielded two MuTx trials (Fig. 2 d – e):

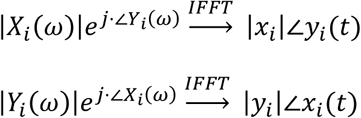

The first of the two MuTx trials, |𝑥_𝑖_|∠𝑦_𝑖_(𝑡), inherited the magnitude information from the CC individual’s 𝑖^th^ trial, but the phase information from the MCC individual’s 𝑖^th^ trial. Conversely, the second MuTx trial, |𝑦_𝑖_|∠𝑥_𝑖_(𝑡), had the magnitude of the MCC individual’s 𝑖^th^ trial but the phase of the CC individual’s 𝑖^th^ trial. Averaging across these trials provided two MuTx ERPs, which we term |𝐶𝐶|∠𝑀𝐶𝐶 and |𝑀𝐶𝐶|∠𝐶𝐶. Here, |𝐶𝐶|∠𝑀𝐶𝐶 is the ERP derived from the trials with the magnitude of CC individuals and the phase of MCC individuals. Conversely, |𝑀𝐶𝐶|∠𝐶𝐶 had the magnitude of MCC individuals and the phase of CC individuals. If a generally lower activation magnitude across trials would be the main driver of the P1 reduction in CC individuals, we would expect |𝐶𝐶|∠𝑀𝐶𝐶 to have lower P1 values than |𝑀𝐶𝐶|∠𝐶𝐶. In contrast, if the P1 wave reduction would be driven mainly by a reduced phase alignment (i.e., coherence) in the CC individuals, then |𝑀𝐶𝐶|∠𝐶𝐶 would exhibit a reduced P1 wave compared to the |𝐶𝐶|∠𝑀𝐶𝐶.

The role of magnitude and phase information in a constellation of sight recovery group and their control participants (i.e., CC vs. MCC or DC vs. MCC) was investigated using the mean of the standardized P1 wave activation (i.e., *z*-score) over the 13 posterior electrodes (120 – 170 ms post-stimulus at electrodes TP9/TP10, CP1/CP2, CP5/CP6, P7/P8, P3/P4, Pz, and O1/O2) as the dependent variable (Sourav et al., 2020).

The *m*ean *p*osterior *P*1 activation, termed MPP1, was investigated in Bayesian linear models of the form MPP1 ∼ magnitude × phase. The factors *magnitude* and *phase* had two levels each, which indicated the source group for the magnitude or phase information, namely *sight recovery* and *control*. Separate models were run for the CC vs. MCC and the DC vs. MDC comparisons.

In addition to this model, we used a reparametrized model with custom contrasts (Schad et al., 2020) to test for three differences of interest. For the CC vs. MCC comparison, the first of these differences tested whether the CC group exhibited a P1 wave reduction (contrast: MCC – CC). The other two contrasts tested the effect of “transplanted” phase on the P1 wave amplitude (contrasts: |CC|∠MCC – CC, and |MCC|∠CC – MCC). Analogous models were run for the DC vs. MDC comparison. In all cases, weak priors were set with the auto_prior() function of the *sjstat* library (Lüdecke, 2018).

### P1 Wave Source Analysis

Average-referenced ERP voltage topographies were converted to a 15,002-point, unconstrained cortical source space using the sLORETA method for EEG inverse modeling (Pascual-Marqui, 2002), implemented in the Brainstorm software package (Tadel et al., 2011). For calculating the forward model used in the sLORETA modeling, we started with the *New York Head* Model with six tissue types: scalp, skull, cerebrospinal fluid, gray matter, white matter, and air cavities (Huang et al., 2016). The forward model was then calculated with the DUNEuro partial differential equation solver package using realistic conductivity values for the tissues and with the electrode locations in the present study (Piastra et al., 2021; Schrader et al., 2021). An identity matrix was used as the noise covariance.

Similar to the P1 wave analysis at the sensor level, group comparisons (CC vs. MCC, DC vs. MDC) of source activity were performed in a posterior cortical region of interest that included the occipital, temporal and parietal regions, excluding the postcentral sulcus and the limbic lobe. To this end, the mean sLORETA activation across the posterior region was extracted, with automatic sign inversion implemented in the Brainstorm package for oppositely oriented sLORETA solutions. The vector norm, i.e. magnitude of the mean sLORETA activation, served as the dependent variable for a Bayesian lognormal model of the form norm_of_mean ∼ Group; weakly informative priors were set for the intercept and groups, respectively *N*(0, 10) and *N*(0, 2.5). For evidence synthesis across experiments, we added a random intercept of participants to take participation in both experiments into account, i.e., norm_of_mean ∼ Group + (1|ParticipantID). Substantial evidence was ascertained when Bayes factors exceeded 3 and when the 95% highest-density interval of the posterior was situated outside of a region of practical equivalence of [-0.1, 0.1] (ROPE test, Makowski et al., 2019).

## Results

### Time-Frequency Analysis

#### Experiment 1

*ΔP_rel_.* The cluster-based permutation test in the whole post-stimulus time-frequency range (latency: 0 – 700 ms, frequency: 2 – 40 Hz) did not reveal any significant difference in *ΔP*_rel_ between the cataract reversal groups and their respective control groups (Fig. 3 a – f); CC vs. MCC, most negative cluster mass (± 95% confidence interval, CI): -369.014 ± 0.006, *p* = .118, most positive cluster mass: 28.399, the parametric threshold for clustering was not exceeded; DC vs. MDC, most negative cluster mass: -241.928 ± 0.009, *p* = .293, most positive cluster mass: 37.595, the parametric threshold for clustering was not exceeded.

**Figure 3.**
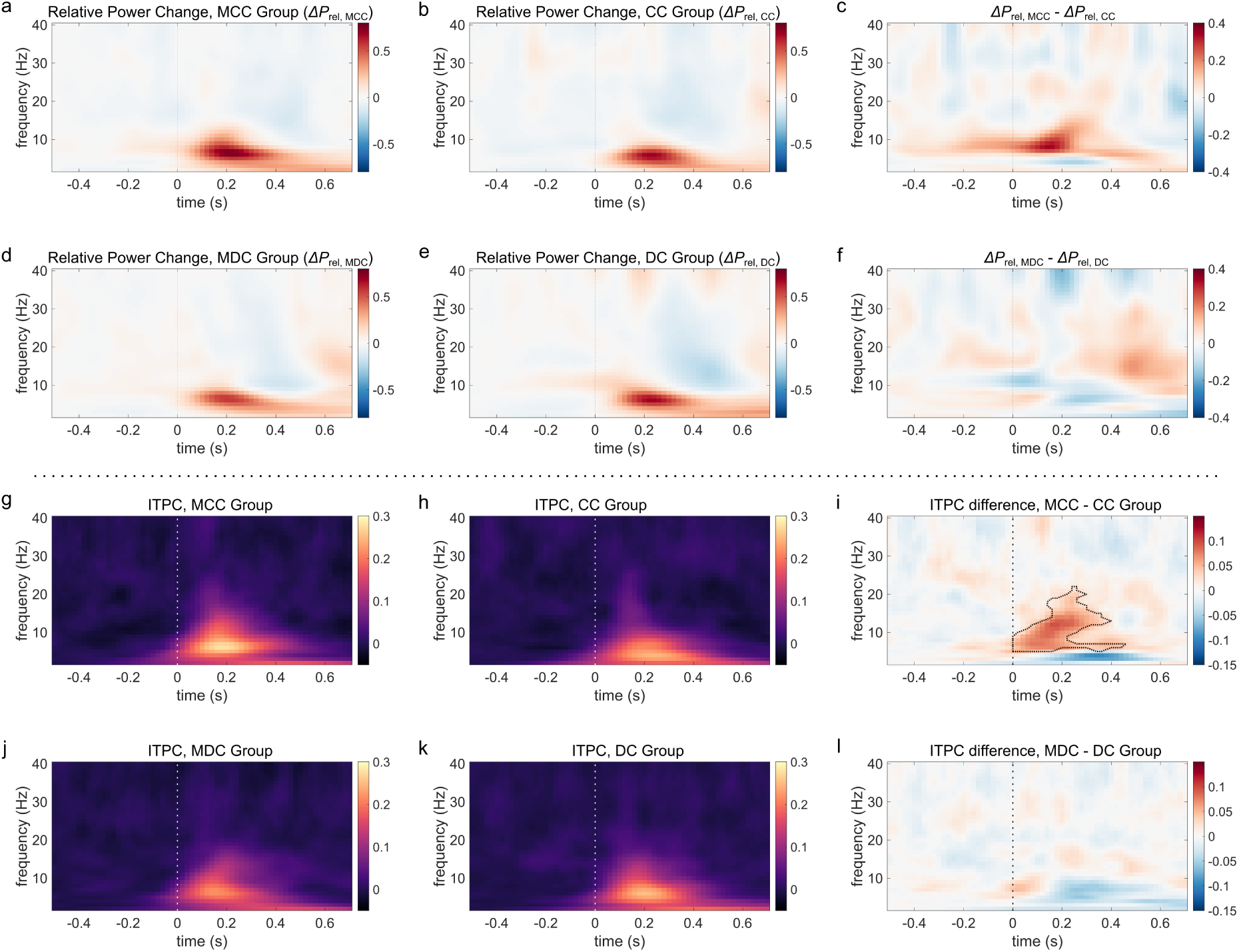
Time-Frequency Analysis of Relative Power Change and Phase Coherence at Posterior Electrodes in Experiment 1. **a** Relative power change (*ΔP*rel, unitless) compared to baseline (-700 to -300 ms) in the matched control group for congenital cataract reversal individuals (MCC). **b** *ΔP*rel in the group of congenital cataract reversal individuals (CC). **c** Relative power change difference between the MCC and the CC group, *ΔP*rel, MCC – *ΔP*rel, CC. **d – f**. Relative power changes in the matched control group for developmental cataract reversal individuals (MDC, *ΔP*rel, MDC), in the group of developmental cataract reversal individuals (DC, *ΔP*rel, DC), and their difference (*ΔP*rel, MDC – *ΔP*rel, DC). **g – i**. Inter-trial phase coherence (ITPC) respectively in the MCC group, the CC group, and their difference. **j – l**. ITPC respectively in the MDC and the DC group, and their difference. For cluster-based permutation tests spanning the time-frequency range of [0 – 0.7 s] × [2 – 40 Hz], the time-frequency extend of a cluster driving a significant result has been outlined. The vertical dotted lines at 0 s indicate stimulus onset. *ΔP*re and ITPC shown in the plots are averaged over the 13 posterior electrodes (TP9/TP10, CP1/CP2, CP5/CP6, P7/P8, P3/P4, Pz, and O1/O2). Note that *ΔP*rel and ITPC have different scales (respectively -0.8 – 0.8 and -0.05 – 0.3). For each variable, the difference between the groups is displayed on a scale that is half of that used for the variable.

Running cluster-based permutation in the “alpha blocking” time-frequency window (time: 0.3 – 0.7 s, frequency: 8 – 12 Hz) as in Bottari et al. (2016), we once more did not find evidence for a difference in *ΔP*_rel_ in the CC vs. MCC analysis (most negative cluster mass = -29.842 ± 0.009, *p* = .304, most positive cluster mass = 0.043, threshold for clustering not exceeded), nor in the DC vs. MDC comparison (most negative cluster mass = -6.959 ± 0.010, *p* = .609, for positive cluster formation no time-frequency point significant at individual level).

*Inter-trial Phase Coherence (ITPC).* In the CC group, significant evidence for a lower ITPC compared to the matched typically sighted control group (MCC) emerged (Fig. 3 g – l); most negative cluster mass (± 95% CI): -405.176 ± 0.003, *p* = .025, most positive cluster mass (± 95% CI): 93.893 ± 0.010, *p* = 0.422. In contrast, between the DC and the MDC group, we did not observe a significant difference in ITPC; most negative cluster mass (± 95% CI): -32.622, the parametric threshold for clustering was not exceeded, most positive cluster mass (± 95% CI): 120.391 ± 0.008, *p* = .226.

#### Experiment 2

*ΔP_rel_.* The cluster-based permutation test in the whole post-stimulus time-frequency range (latency: 0 – 700 ms, frequency: 2 – 40 Hz) once again did not find any significant difference in *ΔP*_rel_ between the CC and the MCC group (Fig. 4 a – f), most negative cluster mass (± 95% confidence interval, CI): -39.6720, the parametric threshold for clustering was not exceeded; most positive cluster mass = 155.132 ± 0.010, *p* = .443. In contrast to the CC vs. MCC group comparison (and in contrast to experiment 1), we found a reduced *ΔP*_rel_ in the DC group compared to the MDC group, most negative cluster mass = -863.053 ± 0.004, *p* = .040; most positive cluster mass = 29.512, parametric threshold for clustering not exceeded.

**Figure 4.**
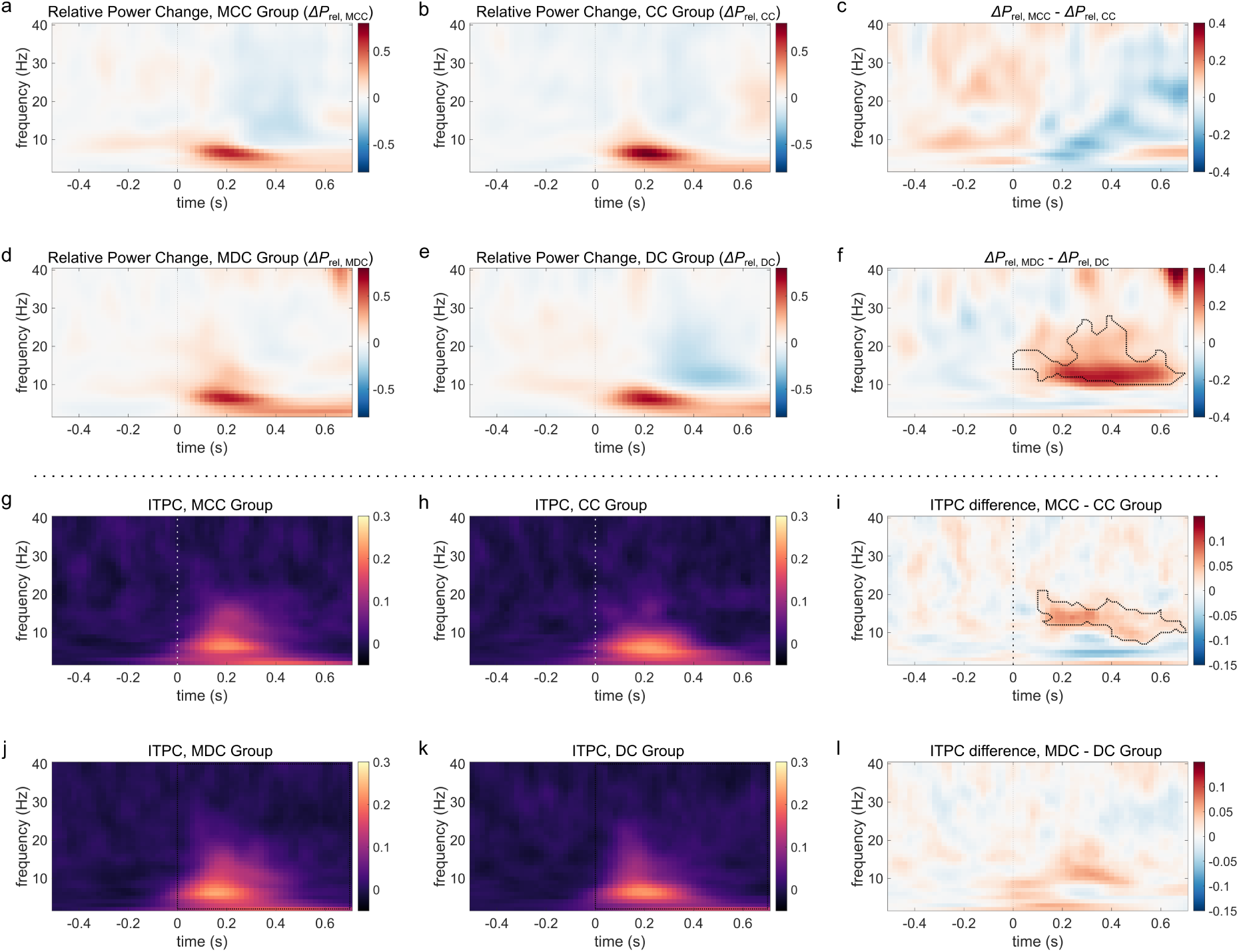
Time-Frequency Analysis of Relative Power Change and Phase Coherence at Posterior Electrodes in Experiment 2. **a** Relative power change (*ΔP*rel, unitless) compared to baseline (-700 to -300 ms) in the matched control group for congenital cataract reversal individuals (MCC). **b** *ΔP*rel in the group of congenital cataract reversal individuals (CC). **c** Relative power change difference between the MCC and the CC group, *ΔP*rel, MCC – *ΔP*rel, CC. **d – f**. Relative power changes in the matched control group for developmental cataract reversal individuals (MDC, *ΔP*rel, MDC), in the group of developmental cataract reversal individuals (DC, *ΔP*rel, DC), and their difference (*ΔP*rel, MDC – *ΔP*rel, DC). **g – i**. Inter-trial phase coherence (ITPC) respectively in the MCC group, the CC group, and their difference. **j – l**. ITPC respectively in the MDC and the DC group, and their difference. For cluster-based permutation tests spanning the time-frequency range of [0 – 0.7 s] × [2 – 40 Hz], the time-frequency extend of a cluster driving a significant result has been outlined. The vertical dotted lines at 0 s indicate stimulus onset. *ΔP*re and ITPC shown in the plots are averaged over the 13 posterior electrodes (TP9/TP10, CP1/CP2, CP5/CP6, P7/P8, P3/P4, Pz, and O1/O2). Note that *ΔP*rel and ITPC have different scales (respectively -0.8 – 0.8 and -0.05 – 0.3). For each variable, the difference between the groups is displayed on a scale that is half of that used for the variable.

**Figure 5.**
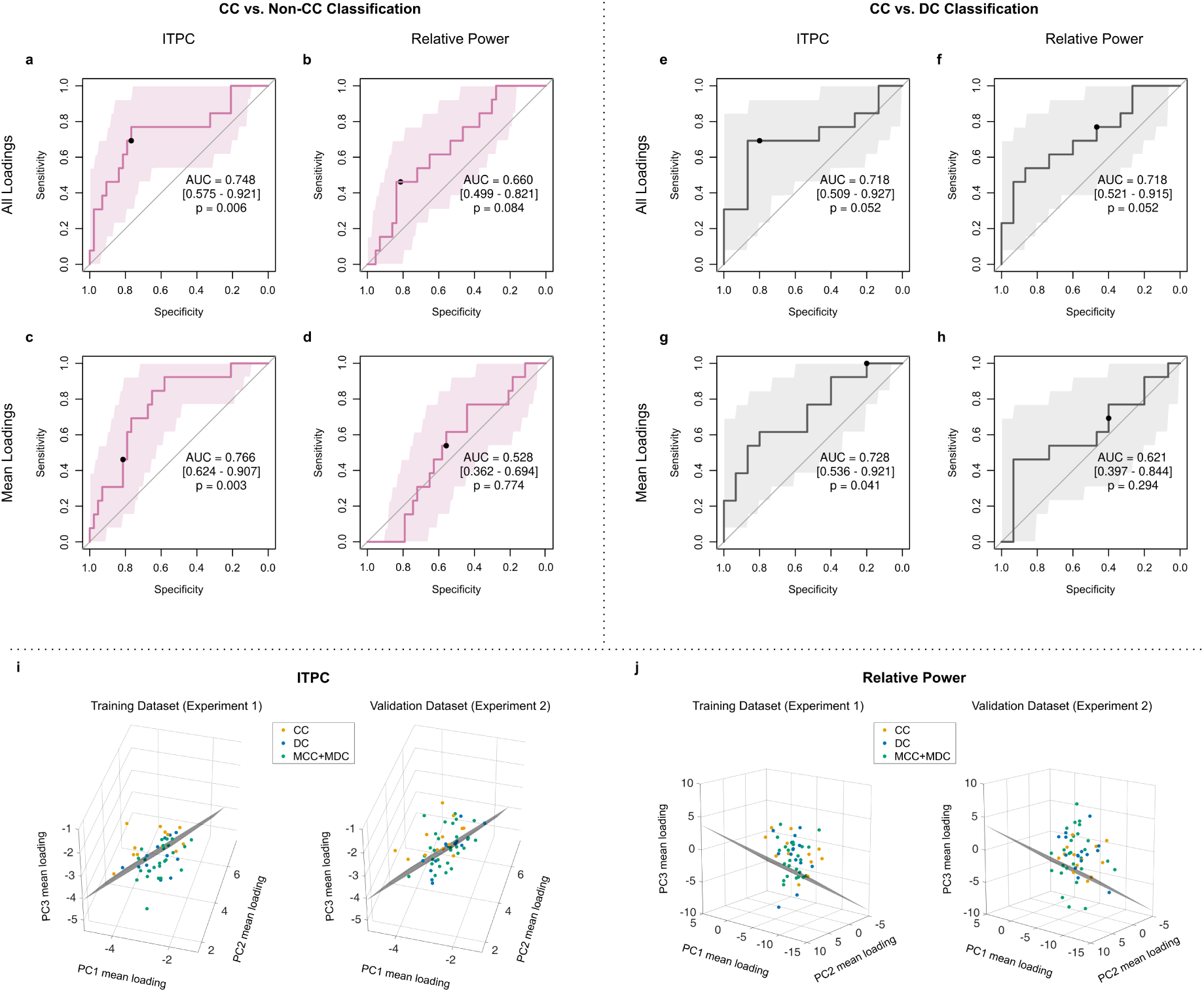
Receiver Operating Characteristic (ROC) Curves for Principal Component Based Classification. **a-d** ROC curves for the classification of CC vs. non-CC individuals based on component loadings for relative power change (*ΔP*rel) and ITPC features at all posterior electrodes (‘All Loadings’) and loadings averaged over posterior electrodes (‘Mean Loadings’). Black dots indicate the optimal classification threshold derived from experiment 1. **e- f** ROC curves for the CC vs. DC classification using the same features. **i** The SVM decision boundary for CC vs. non-CC classification based on ITPC component loadings averaged over posterior electrodes (i.e., 3 features) on the training and validation dataset. **j** The SVM decision boundary for CC vs. Non-CC classification based on component loadings averaged over posterior electrodes for *ΔP*rel on the training and validation dataset.

Employing cluster-based permutation in the “alpha blocking” time-frequency window (time: 0.3 – 0.7 s, frequency: 8 – 12 Hz) as in Bottari et al. (2016), as in experiment 1, no evidence of a difference in *ΔP*_rel_ emerged between the CC and the MCC group (for negative cluster formation, no time-frequency point significant at individual level; most positive cluster mass = 40.989 ± 0.008, *p* = .197).

*Inter-trial Phase Coherence (ITPC).* In the CC group, significant evidence for a lower ITPC compared to the matched sighted control group (MCC) once again was found in experiment 2 (Fig. 4 g – l); most negative cluster mass (± 95% CI): -233.112 ± 0.003, *p* = .031, most positive cluster mass = 28.907, parametric threshold for clustering not exceeded. Similar as in experiment 1, we did not observe a significant difference in ITPC between the DC and the MDC groups; most negative cluster mass (± 95% CI) = -106.505 ± 0.009, *p* = .266; most positive cluster mass = 28.672, parametric threshold for clustering not exceeded.

#### Classification Based on Time-Frequency Features

In addition to the cluster-based permutation tests for group differences in spectral dynamics, we investigated whether CC individuals could be classified based on biomarkers in the time-frequency domain. To prevent overfitting in the high-dimensional spectral feature space, the classifications were performed based on the loadings of the first three principal components at posterior electrodes (39 features), as well as based on the average loadings across the posterior electrodes (3 features). In total, 8 classifications were performed: 2 classification types (CC vs. non-CC, and CC vs. DC) × 2 feature classes (*ΔP*_rel_ and ITPC) × 2 loading types (loadings at all posterior electrodes and loadings averaged over posterior electrodes). All classifiers were trained on data from experiment 1 and validated on data from experiment 2. Validation results are all displayed in Table 5, and the corresponding ROC curves are depicted in Fig. 6, panels a – h. Additionally, the SVM decision boundaries for classification based on ITPC- and power-based component loadings averaged across posterior electrodes are visualized in Fig. 6, panels i – j in the three-dimensional feature space.

**Figure 6.**
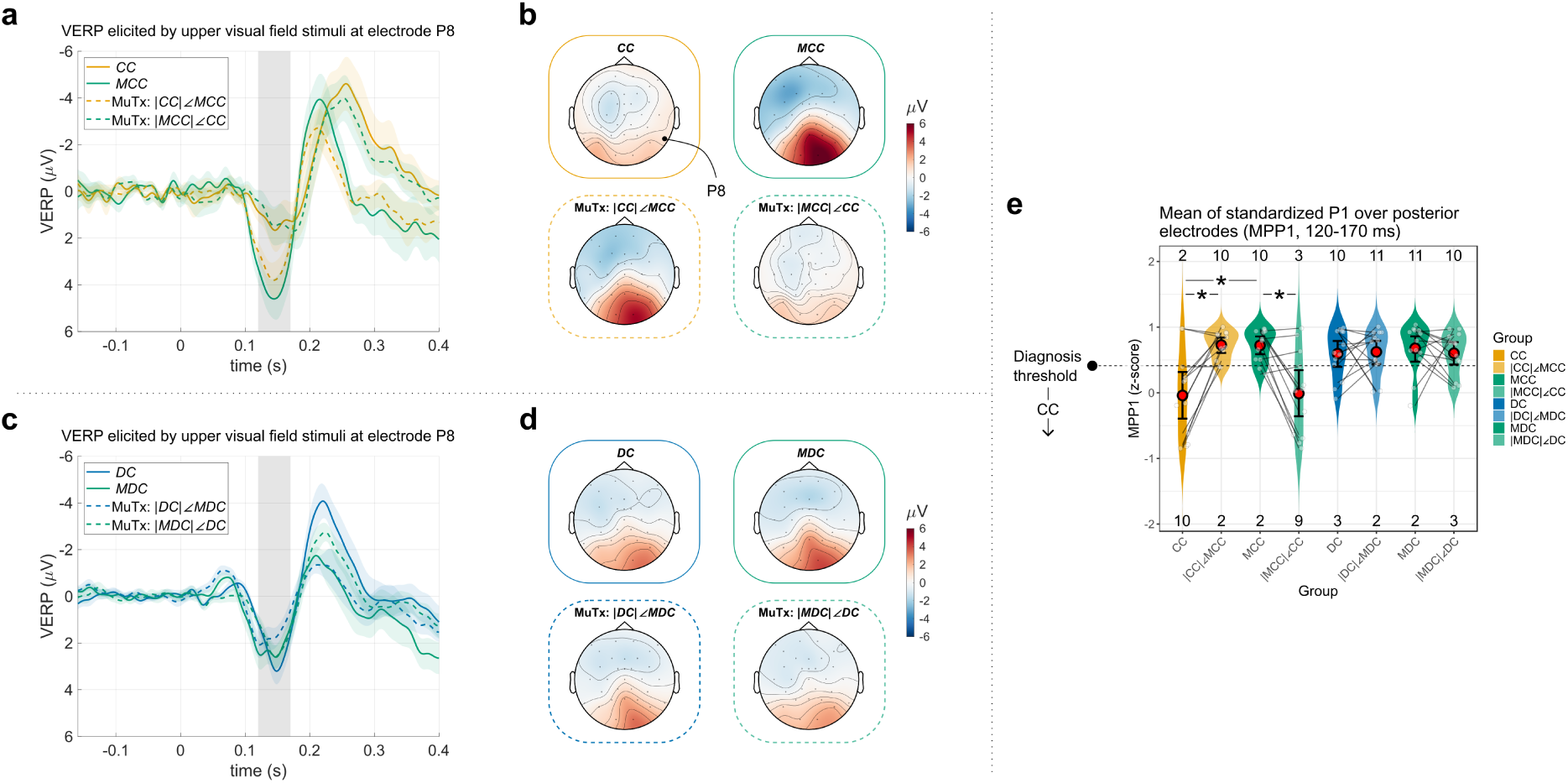
P1 waves of the original groups vs. the mutual magnitude-phase transplant (MuTx) groups in Experiment 1. **a** Grand average event-related potentials (ERP) at the posterior electrode P8 for the sight recovered bilateral congenital cataract group (CC), their matched control group (MCC), and the MuTx groups with the magnitude (|◌|) and phase (∠◌) information swapped, i.e. |CC|∠MCC and |MCC|∠CC. The color-coding is according to the group’s magnitude information. Ribbons indicate the standard error of the mean. The gray bar indicates the P1 wave time range (120 – 170 ms). **b** Topographic plot of the grand average P1 wave for the CC, MCC, |CC|∠MCC, and |MCC|∠CC groups, averaged between 120 – 170 ms after stimulus presentation. **c** Grand average ERPs at the posterior electrode P8 for the developmental cataract reversal group (DC), their matched normally sighted control group (MDC), and the MuTx groups |DC|∠MDC and |MDC|∠DC. **d** Topographic plot of the grand average P1 wave for the DC, MDC, |DC|∠MDC, and |MDC|∠DC groups, averaged between 120 – 170 ms after stimulus presentation. **e** Violin plot of the means of the standardized P1calculated over the posterior electrodes. The mean posterior P1 value (MPP1) had been validated as a retrospective biomarker of congenital visual deprivation (Sourav et al., 2020). The threshold for diagnosing a CC individual is shown as a horizontal dotted line, and the number of individuals above and below the diagnosis line is indicated near the top and bottom border. Red circles denote the means, and the error bars indicate 95% bootstrapped confidence intervals for the means. Asterisks indicate substantial evidence (or above). Individual data points were jittered horizontally for readability.

**Table 5.**
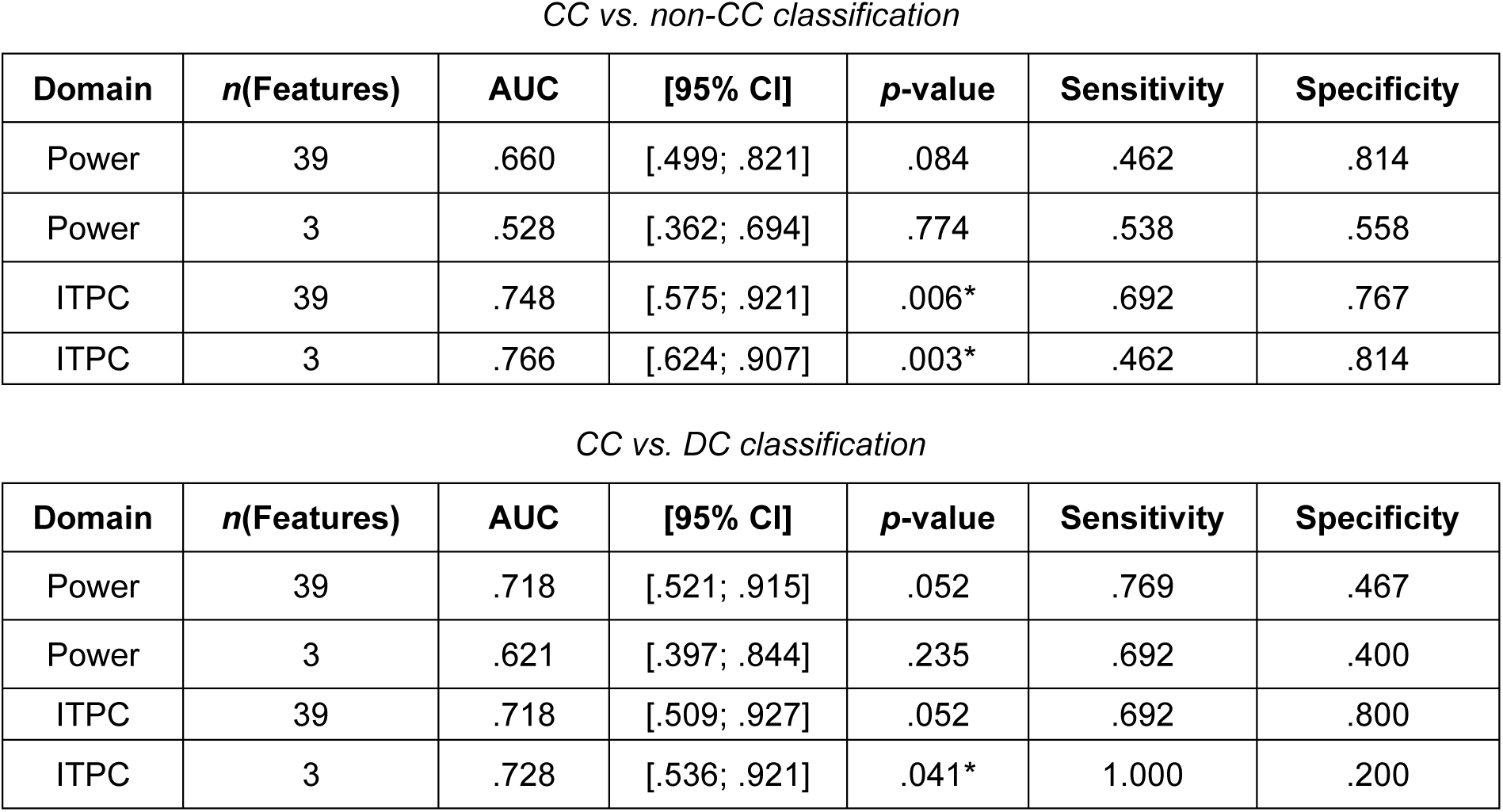
Principal Component Based Classification.

Classification of CC participants from non-CC participants (including DC and normally sighted control individuals) in experiment 2 achieved statistically significant performance based on ITPC component loadings at all posterior electrodes (AUC = .748, 95% CI = [.575 – .921], p =.006), but not based on the component loadings derived from *ΔP*_rel_ (AUC = .660, 95% CI = [.499 – .821], p = .084). Interestingly, a robust performance was observed as well when classification was based on the ITPC component loadings averaged over all posterior electrodes, i.e., based on 3 instead of 39 features (AUC = .766, 95% CI = [.624– .907], p =.003). In contrast, *ΔP*_rel_-based classification performance dropped close to chance level when classifying employed average loadings (AUC = .528, 95% CI = [.362– .694], p = .774). The visualization of the corresponding three-dimensional SVM decision boundaries demonstrates that only the phase-based group differences generalize from the training (experiment 1) to the validation dataset (experiment 2, see Fig. 6 i – j). The optimal thresholds derived from experiment 1 translated to experiment 2 to a moderate degree (Power et al., 2013).

#### CC vs. DC Classification

Classification of the CC from the DC individuals in the validation experiment 2 was not statistically significant using either the ITPC component loadings (AUC = .718, 95% CI = [.509 – .927], *p* = .052) or the *ΔP*_rel_ component loadings at all posterior electrodes (AUC = .718, 95% CI = [.521 – .915], *p* = .052). However, when averaging the component loadings across the posterior electrodes, ITPC information successfully classified the CC from the DC individuals (AUC = .728, 95% CI = [.536 – .921], *p* = .041). In contrast, using average component loadings for *ΔP*_rel_, the classification performance remained close to chance level (AUC = .621, 95% CI = [.397 – .844], *p* = .294).

#### Phase Information is the Main Driver of the P1 Wave Reduction After Sight Recovery Following Transient Congenital Blindness

In a novel application, we exchanged the magnitude and phase information between each sight-recovery individual and their control participant. ERPs were derived from the trials of the mutual magnitude-phase transplanted participants (MuTx) and were systematically compared to those of the original participants (Fig. 6 – 7). We reasoned that if phase information were the main driving factor behind the P1 wave reduction observed in the CC group, the MuTx group comprising the magnitude information of the MCC individuals but the phase information of the CC individuals (denoted |MCC|∠CC) would exhibit a P1 wave reduction. Conversely, the MuTx group consisting of chimeric participants with the magnitude information from the CC individuals but the phase information from the MCC individuals (denoted |CC|∠MCC) would be expected to exhibit typical P1 waves. This was indeed the case: in experiment 1, the grand average visual ERPs qualitatively resembled the group whose phase was used to derive the visual ERP (Fig. 6a). This held true too for the P1 wave topography: in the |CC|∠MCC group’s

P1 wave topography resembled that of the MCC group, and the |MCC|∠CC group’s topography resembled that of the CC group (Fig. 6b). Finally, the Bayesian mixed model investigating the effect of oscillatory magnitude and phase indicated that the mean of the standardized P1 values over posterior electrodes (MPP1) depended on the phase, but not magnitude information content for the CC, MCC, |CC|∠MCC, and |MCC|∠CC constellation; there was a decisive main effect of phase, *β*_phase_ [95% credible interval] = 0.37 [0.23, 0.52], BF_10_ = 1478.71, positive ROPE test. In contrast, there was no substantial evidence for the role of magnitude influencing the MPP1 values in this constellation, *β*_mag_ = 0.01 [-0.14, 0.15], BF_10_ = 0.047, and no substantial interaction between the factors magnitude and phase, *β*_mag × phase_ = -0.01 [-0.15, 0.14], BF_10_ = 0.048.

The same model equipped with custom contrasts provided very strong evidence that the MPP1 values of the CC group was lower than the MPP1 values of the MCC group, *β* = -0.74 [-1.13, - 0.34], BF_10_ = 56.123, positive ROPE test. In addition, the |CC|∠MCC group had a higher MPP1 value than the CC group, *β* = 0.75 [0.34, 1.15], BF_10_ = 71.263, positive ROPE test. Finally, there was very strong evidence that |MCC|∠CC group’s MPP1 values were lower compared to the MPP1 values of the MCC group, *β* = -0.71 [-1.11, -0.31], BF_10_ = 48.557, positive ROPE test, indicating that difference in phase information likely caused the P1 wave attenuation in the CC group.

These results were replicated in the validation dataset from experiment 2, both qualitatively (Fig. 7 a – d) as well as quantitatively (Fig. 7e). Specifically, a Bayesian mixed model once again indicated that the MPP1 values for the CC, MCC, |CC|∠MCC, and |MCC|∠CC constellation depended on the phase but not magnitude information; there was a decisive main effect of phase, *β*_phase_ [95% credible interval] = 0.27 [0.14, 0.40], BF_10_ = 117.02, positive ROPE test. In contrast, there was no substantial evidence for the role of magnitude influencing the MPP1 values in this constellation, *β*_mag_ = 0.06 [-0.07, 0.19], BF_10_ = 0.075, and no substantial interaction between the factors magnitude and phase, *β*_mag × phase_ = -0.01 [-0.14, 0.12], BF_10_ = 0.049.

**Figure 7.**
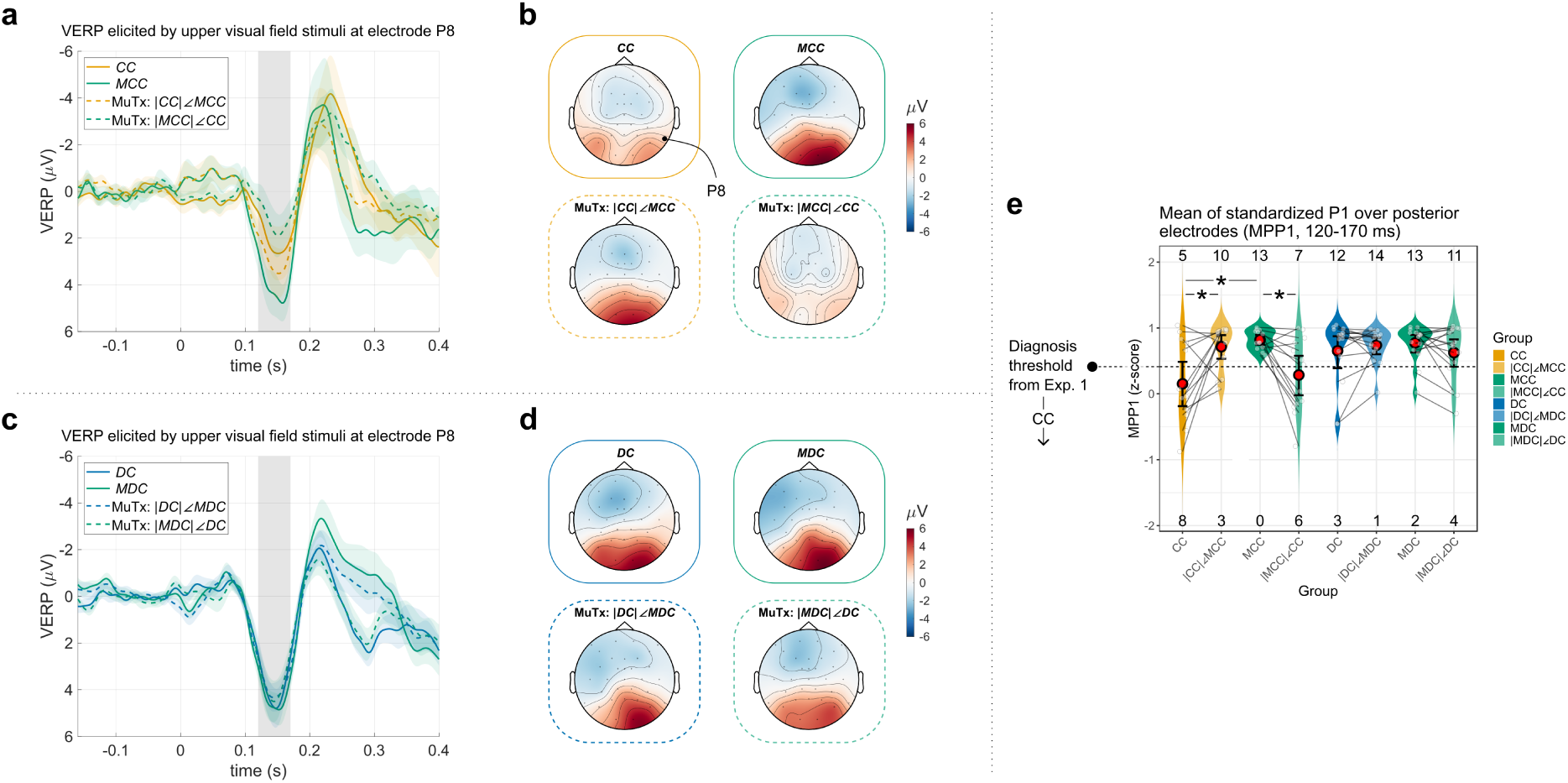
P1 waves of the original groups vs. the mutual magnitude-phase transplant (MuTx) groups in Experiment 2 (Validation Dataset) **a** Grand average visual event-related potentials (VERP) at the posterior electrode P8 for the sight recovered bilateral congenital cataract group (CC), their matched control group (MCC), and the MuTx groups with the magnitude (|◌|) and phase (∠◌) information swapped, i.e. |CC|∠MCC and |MCC|∠CC. The color-coding is according to the group’s magnitude information. Ribbons indicate the standard error of the mean. The gray bar indicates the P1 wave time range (120 – 170 ms). **b** Topographic plot of the grand average P1 wave for the CC, MCC, |CC|∠MCC, and |MCC|∠CC groups, averaged between 120 – 170 ms after stimulus presentation. **c** Grand average VERPs at the posterior electrode P8 for the typically sighted recovered bilateral developmental cataract group (DC), their matched control group (MDC), and the MuTx groups |DC|∠MDC and |MDC|∠DC. **d** Topographic plot of the grand average P1 wave for the DC, MDC, |DC|∠MDC, and |MDC|∠DC groups, averaged between 120 – 170 ms after stimulus presentation. **e** Violin plot of the mean of the standardized P1 topography, with the mean calculated over the posterior electrodes. The mean posterior P1 value (MPP1) was developed and validated as a biomarker based on the present experiments (Sourav et al., 2020). The threshold for diagnosing a CC individual (from experiment 1) is shown as a horizontal dotted line, and the number of individuals above and below the diagnosis line is indicated near the top and bottom border. Red circles denote the means, and the error bars indicate 95% bootstrapped confidence intervals for the means. Asterisks indicate substantial evidence (or above). Individual data points were jittered horizontally for readability.

The same model equipped with custom contrasts provided very strong evidence that the MPP1 values of the CC group were lower than those of the MCC group, *β* = -0.65 [-1.01, -0.28], BF_10_ = 39.909, positive ROPE test. In addition, there was substantial evidence that the |CC|∠MCC group had higher MPP1 values than the CC group, *β* = 0.55 [0.18, 0.91], BF_10_ = 9.893, positive ROPE test. Finally, there was substantial evidence that the |MCC|∠CC group’s MPP1 values were lower compared to those of the MCC group, *β* = -0.52 [-0.88, -0.15], BF_10_ = 6.836, positive ROPE test.

When we tested the DC, MDC, |DC|∠MDC, and |MDC|∠DC group constellation, we found no evidence in any of the two experiments that the MPP1 values substantially differed between the groups (all non-intercept *β*s had BF_10_ < 1, no positive ROPE tests). Unsurprisingly, there was no substantial evidence that the MPP1 depended on phase or the magnitude information in this constellation (all non-intercept BF_10_ < 1, no positive ROPE tests).

#### Source Localization of the P1 wave

In experiment 1, the Bayesian lognormal model indicated that the magnitude of mean posterior sLORETA activation substantially differed between the CC and the MCC groups (Fig. 8), *β*_CC- MCC_ [95% CrI] = -1.03 [-1.76, -0.30], BF_10_ = 6.19, positive ROPE test. In contrast, no substantial difference in magnitude of mean posterior sLORETA activations was observed when comparing the DC vs. the MDC group; *β*_DC-MDC_ = -0.11 [-0.63, 0.41], BF_10_ = 0.111, negative ROPE test (3.06 % of posterior distribution in the ROPE region).

**Figure 8.**
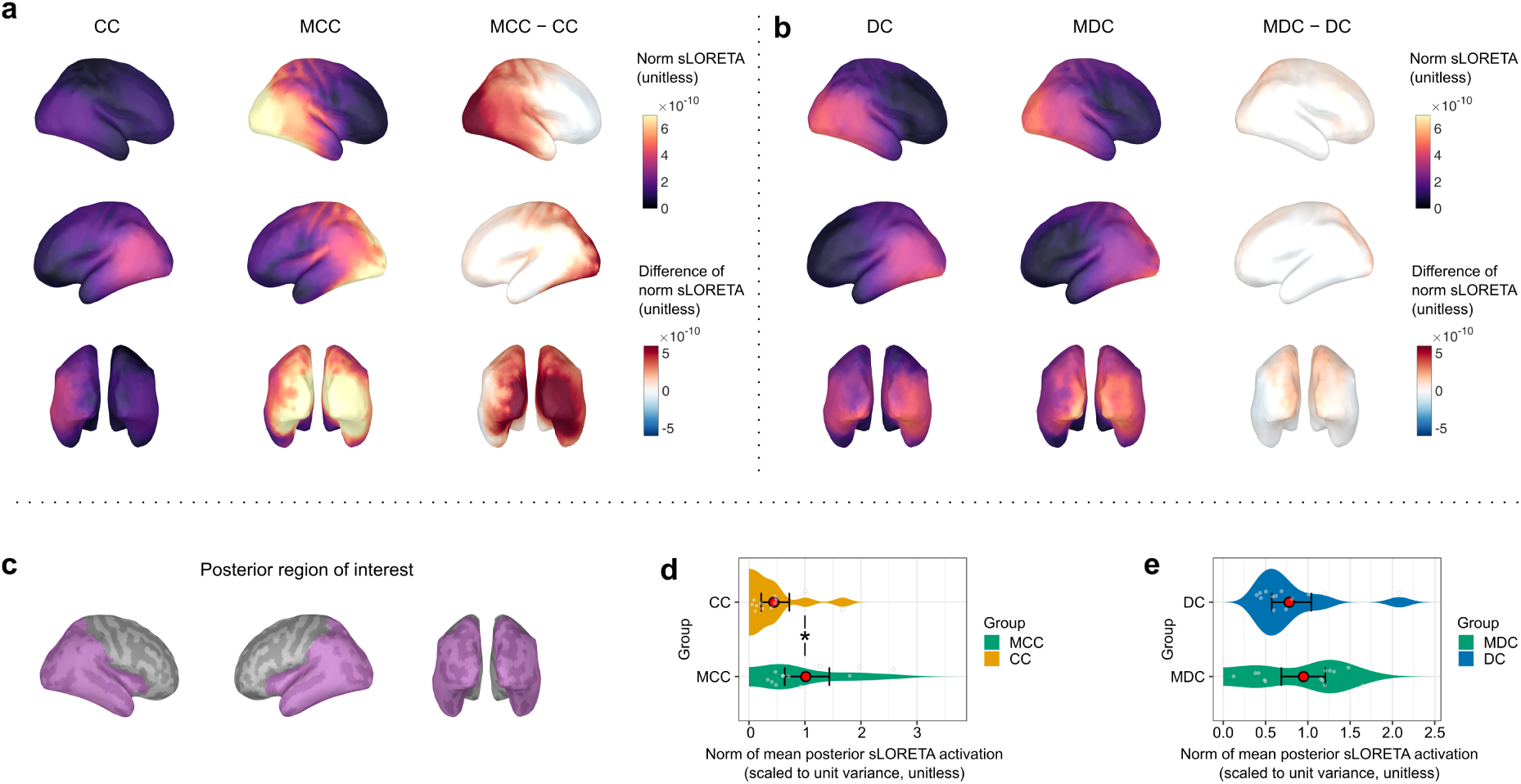
Source Localization of the P1 Wave in Experiment 1. **a** Magnitude (vector norm) of the grand average source localization solution for the P1 wave (mean over 120 – 170 ms post-stimulus) obtained with the sLORETA method (unitless; Pascual-Marqui, 2002; Tadel et al., 2011) respectively for the CC and the MCC groups (left and center columns), and the MCC – CC group difference of grand average norms (right column). **b** Magnitude of the grand average sLORETA solution for the P1 wave (mean over 120 – 170 ms post-stimulus) respectively for the DC and the MDC groups (left and center columns), and the MDC – DC group difference of grand average norms (right column). **c** Posterior region of interest (ROI) for testing group differences, which included the occipital, temporal and parietal regions excluding the postcentral sulcus and the limbic lobe. **d** Violin plot of the magnitude of the mean sLORETA activation in the posterior ROI for the CC and the MCC group (scaled to unit variance). **e** Violin plot of the magnitude of the mean sLORETA activation in the posterior ROI for the DC and the MDC group (scaled to unit variance). Asterisk (*) indicates substantial evidence for a group difference (Bayes factor > 3 and positive ROPE test, lognormal model).

In experiment 2, we did not find any substantial evidence that the magnitude of mean posterior sLORETA activation substantially differed between the CC and the MCC groups (see Fig. 9), *β*_CC-MCC_ [95% CrI] = -0.58 [-1.19, -0.03], BF_10_ = 0.772, negative ROPE test (0.46% of posterior distribution in the ROPE region). No substantial difference in magnitude of mean posterior sLORETA activations was observed when comparing the DC vs. the MDC groups as well; *β*_DC- MDC_ = -0.11 [-0.63, 0.41], BF_10_ = 0.111, negative ROPE test (3.06 % of posterior distribution in the ROPE region).

**Figure 9.**
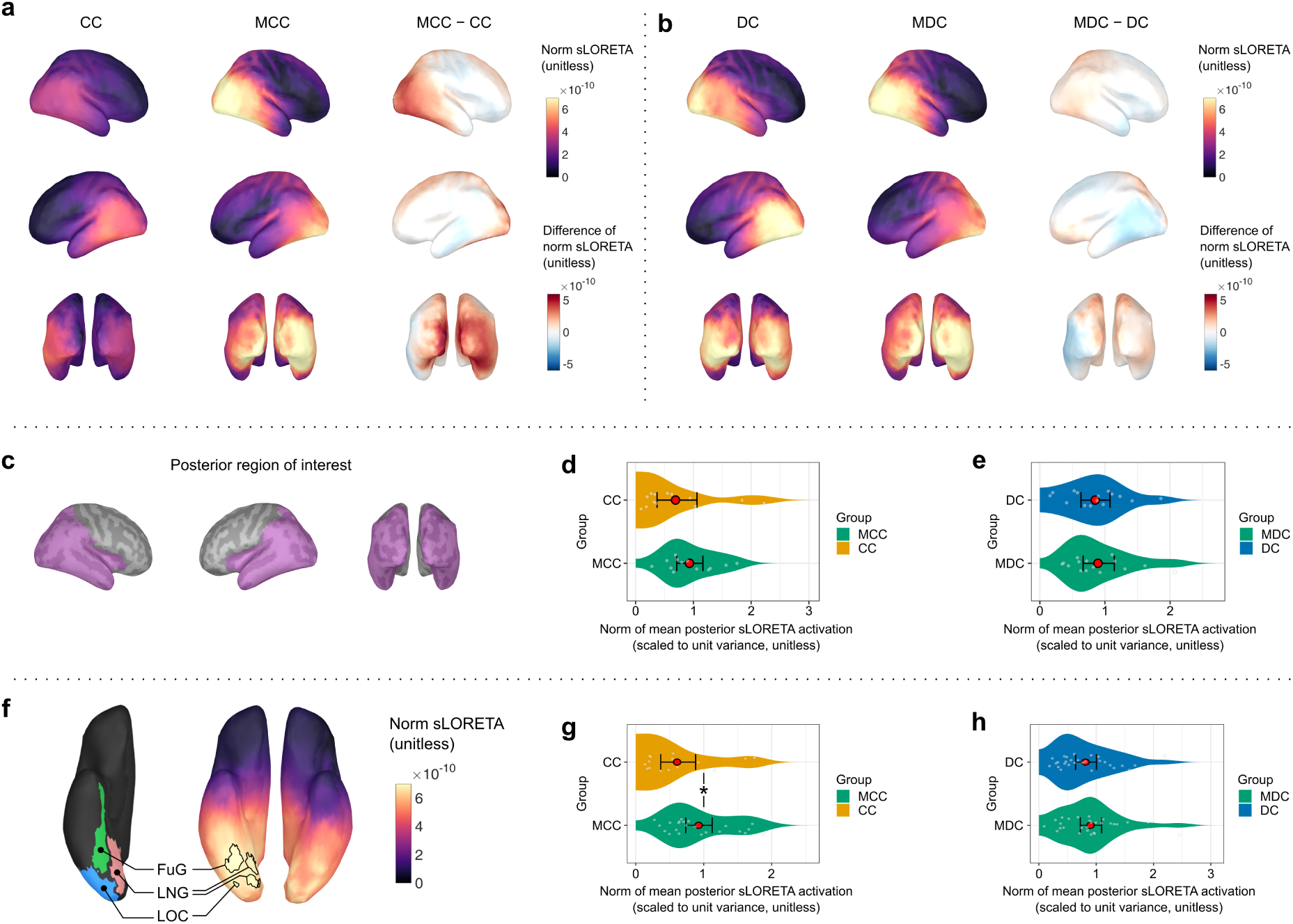
Source Localization of the P1 Wave in Experiment 2 and Evidence Synthesis Across Experiments 1 and 2. **a** Magnitude (vector norm) of the grand average source localization solution for the P1 wave (mean over 120 – 170 ms post-stimulus) obtained with the sLORETA method in experiment 2 (unitless; Pascual-Marqui, 2002; Tadel et al., 2011) respectively for the CC and the MCC groups (left and center columns), and the MCC – CC group difference of grand average norms (right column). **b** Magnitude of the grand average sLORETA solution for the P1 wave (mean over 120 – 170 ms post-stimulus) respectively for the DC and the MDC groups in experiment 2 (left and center columns), and the MDC – DC group difference of grand average norms (right column). **c** Posterior region of interest (ROI) for testing group differences, which included the occipital, temporal and parietal regions excluding the postcentral sulcus and the limbic lobe. **d** Violin plot of the magnitude of the mean sLORETA activation in the posterior ROI for the CC and the MCC group in experiment 2 (scaled to unit variance). **e** Violin plot of the magnitude of the mean sLORETA activation in the posterior ROI for the DC and the MDC group in experiment 2 (scaled to unit variance). **f** Grand average source localization solution across all participants and groups in experiment 1 and experiment 2. Black borders indicate the loci of the highest 5% sLORETA magnitude. Relevant cortical regions from the Desikan-Killiany atlas are depicted on the left side of the sLORETA solution; FuG: Fusiform gyrus, LNG: Lingual gyrus, LOC: Lateral occipital cortex. **g** Violin plot of the magnitude of the mean sLORETA activation in the posterior ROI for the CC and the MCC group across experiment 1 and experiment 2, scaled to unit variance. All unique participants are depicted, for participants who took part in both experiments, mean values are presented. **h** Violin plot of the magnitude of the mean sLORETA activation in the posterior ROI for the DC and the MDC group across experiment 1 and experiment 2, scaled to unit variance. All unique participants are depicted, for participants who took part in both experiments, mean values are presented. Asterisk (*) indicates substantial evidence for a group difference (Bayes factor > 3 and positive ROPE test, lognormal model)

The Bayesian models, being parametric, allowed evidence synthesis across the experiments, which on the one hand led to a more stable parameter estimation and on the other hand permitted reconciling partially different result patterns for the CC vs. MCC group difference across the experiments. Combining evidence across both experiments with Bayesian lognormal mixed models, we found substantial evidence for a group difference in the mean posterior sLORETA activation strength between the CC and the MCC group, *β*_CC-MCC_ [95% CrI] = -0.77 [-1.26, -0.27], BF_10_ = 9.01, positive ROPE test (Fig. 9). In contrast, once again no substantial difference in the norm of mean posterior sLORETA activations emerged when comparing the DC vs. the MDC group; *β*_DC-MDC_ = -0.10 [-0.46 0.27], BF_10_ = 0.085, negative ROPE test (3.97 % of posterior distribution in ROPE region).

The grand average sLORETA map across all participants and both experiments indicated posterior regions as the source of the P1 wave (Fig. 9). The maximum 5% regions of this activation were located in the right ventral extrastriate visual cortical regions, including the fusiform gyrus (FuG), lingual gyrus (LNG) and lateral occipital cortex (LOC) according to the Desikan-Killany atlas (Desikan et al., 2006).

## Discussion

Optimal functioning of neural circuits requires appropriate neural activation strength as well as precisely timed excitation and inhibition. Since sufficiently temporally resolved neural timing information is hard to access with brain imaging methods (e.g., fMRI), and as activation strength and timing information is intermingled in trial-averaged electrophysiological measures such as those derived from EEG/MEG, how neural circuit timing depends on early experience is still an unanswered question. A recent study involving ferrets has suggested that particularly temporal processing stability in the visual cortex may depend on visual input after birth (Trägenap et al., 2025). However, in non-human animal research the degree of recovery after visual deprivation and subsequent sight restoration is difficult to investigate. Thus, it is currently unknown whether a sensitive period for temporally tuned neural circuit processing exists; sensitive periods are defined by incomplete recovery after sensory restitution (Kundsen, 2004).

A transient phase of congenital visual deprivation in humans has been found to permanently alter visual cortical structure and function (Le Grand et al., 2003; Röder et al., 2021; Hölig et al., 2023; Pedersini et al., 2023; Sourav et al., 2024). Of specific relevance is the earliest electrophysiological maker of higher-order, extrastriate visual cortical activity, the P1 wave, which peaks at around 150 ms after the visual stimulation onset (Di Russo et al., 2002). The P1 wave was found to be markedly attenuated in sight-recovery individuals with a history of dense, bilateral congenital cataracts (Sourav et al., 2020). Yet, to the best of our knowledge, no study has systematically examined the trial-to-trial temporal consistency of cortical oscillations in individuals with reversed congenital blindness to disentangle to role of cortical activation strength vs. impaired temporal processing stability as a neural mechanism of their extrastriate visual processing impairment. A dissociation of both aspects of neural processing is crucial for the understanding of the precise mechanisms of experience-dependent brain development.

The present study investigated group differences in task-related oscillatory activation strength and timing stability in individuals with reversed congenital cataracts (CC), individuals with reversed developmental, i.e., late-onset cataracts (DC), and matched normally sighted controls (SC). Three methodological approaches were implemented: First, in time-frequency representations of cortical activity during the processing of simple visual stimuli, the across-trial consistency of visual processing, indexed by the inter-trial phase coherence, was significantly attenuated in the CC group compared to their matched normally sighted controls (MCC group). In contrast, no differences in relative power change from baseline between these groups was observed. Second, further supporting these findings, the CC individuals could be classified from non-CC individuals (i.e., a combined group of sighted controls as well as DC individuals) only by using phase information but not by using oscillatory power information. Third, through a novel approach that we term *mutual magnitude-phase transplants,* we swapped the oscillatory activation magnitude and phase information between each CC individual and their matched sighted controls. This analysis revealed that combining the phase information of the CC individuals with the magnitude information of the MCC individuals, but not vice versa, yielded the attenuation of the extrastriate P1 wave in visual event-related potentials (ERPs) which is typical for the CC group. The convergent evidence from all three methodological approaches was replicated in a second experiment. Thus, task-related oscillatory timing information, but not oscillatory strength, appears to critically depend on early experience and therefore seems to be a key mechanism of sensitive period plasticity.

Between the DC individuals and their matched control group (MDC), cluster-based permutation tests did not reveal significant temporal stability differences in any experiment, indicating that the decrease in task-related oscillatory phase synchronization in the CC vs. MCC group comparison is linked to visual deprivation during an early sensitive period. This selective impairment for CC individuals excludes alternative explanations such as unspecific effects related to cataract surgery. Additionally, phase, but not power information averaged over the posterior electrodes allowed us to classify CC from DC individuals. Lastly, *mutual magnitude-phase transplant* analyses revealed no effect of exchanging the magnitude and phase information between the DC and the MDC individuals, confirming that temporal processing impairments were specific to the CC group.

Compared to striate cortical processing, extrastriate cortical functions have been reported to exhibit higher vulnerability to a period of transient congenital visual deprivation (McKyton et al., 2015; Sourav et al., 2018, 2020; Pitchaimuthu et al., 2021). Consistent with this line of evidence, we previously found a robustly reduced P1 wave in CC individuals’ visual ERPs (Sourav et al., 2020), which has been hypothesized to originate in extrastriate cortical areas (Di Russo et al., 2002; Miller et al., 2015). Moreover, this strong and reliable reduction of the P1 wave in sight recovery individuals allowed a robust retrospective diagnosis of a congenital vs. developmental etiology years after sight restoration (Sourav et al., 2020). In the present study, source localization of the P1 wave indicated areas of maximum activation in extrastriate cortical areas in the right (ipsilateral) hemisphere, including the fusiform gyrus, lingual gyrus and lateral occipital cortex. These results are by and large consistent with previous EEG/fMRI work (Di Russo et al., 2002). Moreover, augmenting the sensor-level findings, evidence synthesis across the two experiments of the present study revealed substantial evidence for a trial-averaged reduction of the P1 wave in the CC group at source level as well.

Phase alignment of alpha oscillatory activity (8 – 12 Hz) has been argued to contribute to the generation of the P1 wave (Klimesch et al., 2007), supported by the findings that alpha phase alignment predicts the P1 wave latency (Gruber et al., 2005), and that alpha phase locking is largest during the P1 wave latency (Klimesch et al., 2004; see Freunberger et al., 2008). Consistent with this evidence, the principal component loading patterns of the inter-trial phase coherence features for classifying CC individuals were predominantly located across the alpha (8 – 12 Hz) and the theta (4 – 8 Hz) range and covered the P1 wave latency. Additionally, significant inter-trial phase coherence differences between the CC and the MCC group in the cluster-based permutation tests were associated with time-frequency clusters that spanned the P1 latency and the alpha frequency range. In contrast to phase alignment across trials, the P1 wave amplitude has been argued to be less affected by oscillatory power (Gruber et al., 2005; Freunberger et al., 2008). In a related visual deprivation model of amblyopia, a reduction of the P1 wave amplitude in the patients’ amblyopic eye compared to the fellow eye was associated with a broader trial-to-trial latency (but not amplitude) distribution (Bankó et al., 2013). Consistent with these lines of evidence, we did not find any significant difference in the post-stimulus relative power change in the CC vs. MCC group comparison. We did not observe a significant classification performance based on relative power change as opposed to inter-trial phase coherence information either. In fact, the *mutual magnitude-phase transplants* analysis in the present study provided compelling evidence of the driving role of phase information behind P1 reduction in the CC group: combining CC individuals’ activation magnitudes with the phase information of typically sighted controls substantially recovered their P1 wave amplitude.

In the present study, we did not observe any relative oscillatory power impairments in the CC group compared to the sighted controls during the processing of simple visual stimuli (gratings) in any of the two experiments. While these results are inconsistent with those of Bottari et al. (2016) in a group of CC individuals (*n* = 12) during biological motion processing, they are corroborated by Ossandón et al (2023): The latter authors have recently reported a spared alpha power modulation in resting state EEG due to eye opening (compared to an eyes closed condition) in CC individuals, despite an overall lower alpha oscillatory power level compared to sighted controls.

Temporal stability, and its converse, temporal variability, is a critical marker of sensory processing related to the alignment of dynamic neural states (Afraimovich et al., 2004; Jones et al., 2007; Rabinovich et al., 2008). The converging results of the two present experiments indicate that a disrupted temporal orchestration of cortical activity (Yusuf et al., 2022), especially in higher sensory areas, and not a general attenuation of cortical oscillation may be the most profound consequence of a period of congenital sensory deprivation. In this context we note that feedforward and feedback connection profiles develop asynchronously, with feedforward connectivity maturing earlier (Burkhalter et al., 1993). Moreover, there is evidence that the elaboration of feedback connectivity more extensively depends on experience (Ibrahim et al., 2021; Röder and Kekunnaya, 2021). Early sensory experience, thus, might be crucial for temporally aligned cortical feedback signaling and in turn for the shaping of feedforward activity (Ibrahim et al., 2021). A recent study that examined the development of the ferret visual cortex strongly supports this account: in ferret kits whose eyes were prematurely opened, visual cortical responses recorded with calcium imaging were found to be similarly strong, yet temporally less stable across trials compared to visually experienced kits (Trägenap et al., 2025). Crucially, a week of lid suture in the ferret kits, starting shortly before the time of natural eye opening, disrupted the typical development of the temporal stability of cortical processing when tested shortly after lid suture removal. The present results, on the one hand, extend these findings in ferrets to human cortical development. On the other hand, the temporal resolution of the Trägenap et al. (2025) study was much lower than that of the EEG recordings we employed: The specific calcium indicator utilized by the authors (GCaMP6s, see Trägenap et al., 2025) had a time constant of over a hundred milliseconds, that is, a temporal resolution two orders of magnitude less precise than EEG which provides millisecond-level temporal resolution (Buzsáki et al., 2012; Jercog et al., 2016). Thus, the results of the present study extend the findings of Trägenap et al. (2025): The disrupted temporal stability of cortical processing, resulting from aberrant early visual experience seems to appear already within 150 ms after visual stimulation onset in markers of extrastriate and downstream visual cortical processing. Crucially, we investigated sight-recovery individuals at least two years after sight restitution surgery. Thus, we are able to report evidence for a sensitive period in humans, that is, for a persistent loss of temporal processing stability in visual neural circuits despite sight restitution and extensive post-surgical visual experience.

In conclusion, we provide to the best of our knowledge the first report that temporal processing stability in visual neural circuits critically depends on early experience in humans. This finding of an impaired visual cortical processing stability in humans after a period of transient congenital (but not developmental) pattern visual deprivation provides strong evidence for the existence of a sensitive period for the development of precisely temporally orchestrated visual cortical processing in humans. Our results suggest that aberrant temporal tuning of the visual cortex, especially in extrastriate and higher-order visual areas, might underlie the numerous persistent visual deficits observed after reversing congenital blindness.

## Acknowledgements

We thank D. Balasubramanian (L V Prasad Eye Institute, Hyderabad, India) for supporting this study. We acknowledge Davide Bottari for helping with data acquisition and for the initial EEG preprocessing pipeline. Seema Banerjee, Larissa Brockmann, Maria Guerreiro, Marlene Hense, Giulia Dormal, Siddhart Srivatsav Rajendran, Lisa Stockleben and Florian Süßer additionally helped with data acquisition in India and/or Germany. The study was funded the German Research Foundation (DFG, Ro 2625/10-1) and by the European Research Council grant ERC-2009-AdG 249425-*CriticalBrainChanges* to B.R.

## Notes

### Competing Interest Statement

The authors have declared no competing interest.

### Summary of Updates

The abstract, significance statement, introduction and discussion sections have been streamlined. A CRediT taxonomy has been added for author contributions. The figures, analyses, and results remain unchanged.

